# Nucleosome context regulates chromatin reader preference

**DOI:** 10.1101/2025.04.29.651129

**Authors:** Matthew R. Marunde, Irina K. Popova, Nathan W. Hall, Anup Vaidya, James R. Bone, Brandon A. Boone, Peter J. Brown, Ryan J. Ezell, Tessa M. Firestone, Harrison A. Fuchs, Elisa Gibson, Zachary B. Gillespie, Susan L. Gloor, Allison R. Hickman, Sarah A. Howard, Natalia Ledo Husby, Victoria T. Hsiung, Andrea L. Johnstone, Laiba F. Khan, Krzysztof Krajewski, Alexander S. Lee, Eileen T. McAnarney, Keith E. Maier, Danielle N. Maryanski, Kelsey E. Noll, Katherine Novitzky, Emily F. Patteson, Keli L. Rodriguez, Julio C. Sanchez, Luis F. Schachner, Catherine E. Smith, Lu Sun, Hailey F. Taylor, Rachel Watson, Hannah E. Willis, Catherine A. Musselman, Bryan J. Venters, Marcus A. Cheek, Matthew J. Meiners, Zu-Wen Sun, Neil L. Kelleher, Martis W. Cowles, Ellen N. Weinzapfel, Michael-Christopher Keogh, Jonathan M. Burg

## Abstract

Chromatin is more than a simple genome packaging system, and instead locally distinguished by histone post-translational modifications (PTMs) that can directly change nucleosome structure and / or be “read” by chromatin-associated proteins to mediate downstream events. An accurate understanding of histone PTM binding preference is vital to explain normal function and pathogenesis, and has revealed multiple therapeutic opportunities. Such studies most often use histone peptides, even though these cannot represent the full regulatory potential of nucleosome context. Here we apply a range of complementary and easily adoptable biochemical and genomic approaches to interrogate fully defined peptide and nucleosome targets with a diversity of mono or multivalent chromatin readers. In the resulting data, nucleosome context consistently refined reader binding, and multivalent engagement was more often regulatory than simply additive. This included abrogating the binding of the Polycomb group L3MBTL1 MBT to histone tails with lower methyl states (me1 or me2 at H3K4, H3K9, H3K27, H3K36 or H4K20); and confirmation that the CBX7 chromodomain and AT-hook-like motif (CD-ATL) tandem act as a functional unit to confer specificity for H3K27me3. Further, *in vitro* nucleosome preferences were confirmed by *in vivo* reader-CUT&RUN genomic mapping. Such data confirms that more representative chromatin substrates provide greater insight to biological mechanism and its disorder in human disease.

## INTRODUCTION

Chromatin is an essential regulator of multiple biological processes, including transcription (1–4), DNA damage repair (5,6), cellular differentiation (7–9) and pathogenesis (10–12). Its basic repeating subunit is the nucleosome: a core histone octamer (two each of H2A, H2B, H3 and H4) wrapped with ~147 bp DNA (13). These structures are highly dynamic, and distinguished by / associated with a compendium of histone variants (*e.g.*, H2A.Z, H3.3, CenH3), post-translational modifications (PTMs; *e.g.,* methylation, acetylation, phosphorylation, ubiquitylation), DNA modifications (*e.g.,* methylation; primarily 5-methyl cytosine), and chromatin-associated proteins (CAPs; including histone modifiers, chromatin remodelers and transcription factors) (14–16).

The ‘histone code’ hypothesis posits that combinatorial histone PTMs act as a molecular language ‘read’ by CAPs to facilitate DNA transactions (17,18). This idea was highly controversial at inception (19–21), but has proven hugely insightful and greatly contributed to epigenetics research (22–24). A multiplicity of ‘reader’ domains have now been identified, and understanding how they interact with and interpret histone PTMs can reveal novel disease mechanisms, biomarkers and therapeutic targets (25). Such studies have been challenged by the dynamic nature of chromatin and (until recently) the limited availability of defined nucleosome substrates (24,26–28). Instead, the field has largely relied on modified histone peptides, where low manufacturing costs and compatibility with high-throughput screening (HTS) platforms have supported reader profiling to hundreds of PTM combinations (29–31). However, peptide array screens are often compromised by high material consumption, low signal-to-background (S/B) and highly variable results (32–34). They also rarely include *post hoc* optimizations, which can be essential to deliver maximal insight (28).

Perhaps the greatest underminer of peptide-based approaches is their inability to inform on CAPs that require nucleosome substrates, including NSD2 (35–37), LSD1 (38–40), DOT1L (41,42), EZH2 (43–45) or PR-DUB (46). Such necessity is often due to obligate multivalent engagement, where the CAP contacts multiple histone PTMs, DNA modifications and/or nucleosome surfaces (27,47–56). As an example, NSD2 lysine methyltransferase contains two PWWP domains that interact with nucleosomal DNA to effectively engage H3K36me2 (37). Histone peptides are thus unable to recapitulate the context to fully characterize NSD2 PWWP domain binding (37) and develop maximally effective domain inhibitors (57,58). CAPs can also contain multiple reader domains that interact with individual histone PTMs *in trans* (*i.e.,* on separate histones), as in the PHIP/BRWD2 Tudor-Bromodomain-Bromodomain (Tudor-BD-BD2) reader triplet with ([H3K4me3K14ac]•[H4K12ac]) nucleosomes (59). Interactions with DNA, the histone tails, and/or the H2A/H2B acidic patch can play essential roles in CAP function, such as to allosterically activate DOT1 lysine methyltransferase after engaging H2BK120ub (50–53). Finally, the positive charge on histone tails can be neutralized, such as by lysine acetylation, to relieve the association with negatively charged nucleosomal DNA and increase target accessibility for reader binding and further modification (27,47,55). These examples underscore the need for reliable nucleosome-based assays for effective CAP interrogation.

To address this, we developed the Captify™ platform (Captify for brevity) using no-wash high-sensitivity Alpha (27,28,46,60–62) or washable multiplexable Luminex (63) technologies for the rapid, sensitive and robust screening of CAPs (the Queries) to fully defined histone peptides and nucleosomes (the Targets). The ability to explore diverse conditions, including reactant concentrations and additives, often revealed multivalent engagement mechanisms and refined specificities on nucleosomes. This was confirmed by orthogonal approaches, including reader-CUT&RUN genomic mapping, indicating that *in vitro* CAP : nucleosome binding data more accurately reflects *in vivo* chromatin interactions.

## RESULTS

### Captify-Alpha has dramatically improved performance over peptide arrays and supports nucleosome studies

The dissection of CAP : chromatin engagement mechanisms has historically relied on isolated reader domains and PTM-defined histone peptides. This reductionist approach has advanced our understanding, but physiological relevance is potentially undermined by both components being removed from higher-order context (*i.e.*, protein complexes and nucleosomes). We previously developed histone peptide arrays (*e.g.,* EpiTriton™) for such research, but observed multiple drawbacks, including high Query protein requirements (up to 800 pmol) and low dynamic range. Most notably, the arrays were nucleosome-incompatible, and thus unsuitable for exploring multivalent engagement – a feature of many CAPs. Consequently, ~80% of tested Queries failed to generate reliable binding data (not shown).

To address these shortcomings, we developed Captify on the high-sensitivity no-wash Alpha platform (64) (hereafter Captify-Alpha). Here, biotinylated substrates (the Targets) are mixed with epitope-tagged CAPs (the Queries) and relative engagement (expressed as EC^rel^) quantified through proximity-dependent signal generation using streptavidin “Donor” and anti-tag “Acceptor” beads (27,60–62) **Suppl. Fig. 1A** and **Suppl. Table 1**). Compared to histone peptide arrays, Captify-Alpha is highly adaptable, HTS-compatible, and provides substantial gains in sensitivity and S/B, while dramatically reducing material consumption. To demonstrate, we examined BRD4 bromodomain 1 (GST-BRD4 BD1; Query) binding to histone peptides by EpiTriton array (2 µM Query; hits defined as > two-fold S/B) and Captify-Alpha (30 nM and 1 nM Query; hits defined as > 100-fold S/B). In both formats, BRD4 BD1 bound H4 acetyl-lysine peptides, preferring those with multiple acetylations (*i.e.,* H4_[1-23]_K5acK8acK12acK16ac; *aka*. H4_[1-23]_tetra^ac^) over singles (31,65) (**Figs 1A-B**). However, the Captify-Alpha approach captured more hits, including all detected by peptide arrays, while using much less Query (up to 2,000-fold) and with a much higher dynamic range (>1,000-fold) (**Figs 1A-B**). Further, these binding partners and their rank-ordering were confirmed by orthogonal TR-FRET (65) (**Suppl. Fig. 1B-C**).

**Fig. 1.**
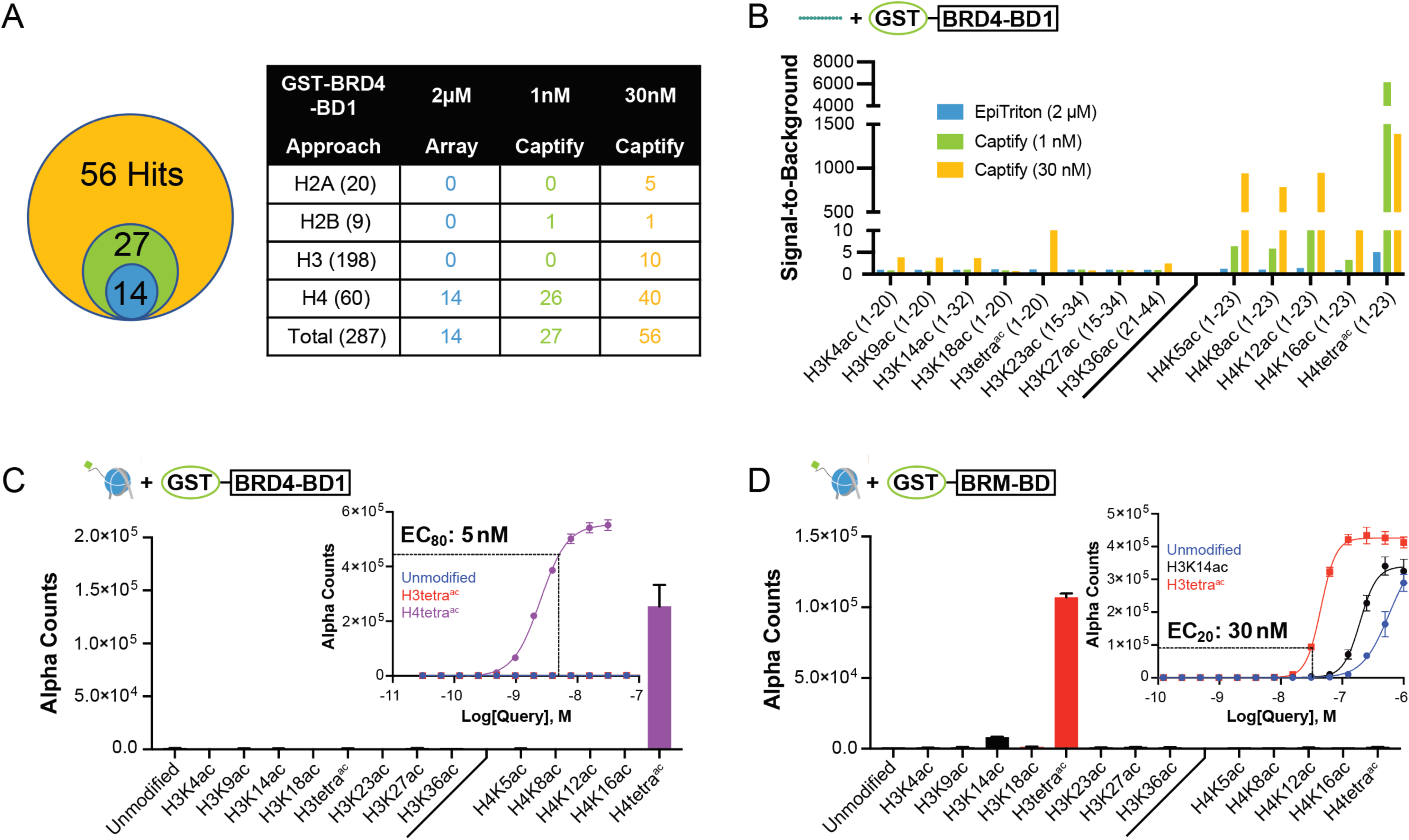
Lysine-acetyl readers show refined PTM specificity in nucleosome *vs.* histone peptide assays. **(A)** GST-BRD4 BD1 (Query, concentrations as noted) interactions in histone-peptide based assays: EpiTriton array and Captify-Alpha. “Hits” are defined by signal-to-background (S/B) >2 for peptide array and >100 for Captify-Alpha. **(B)** GST-BRD4 BD1 S/B in EpiTriton (2 μM) and Captify-Alpha (1 nM or 30 nM) for select H3 and H4 acetyl peptide Targets. Each assay identifies the bromodomain preferentially interacting with various forms of H4 tail acetyl. **(C)** *[Inset]* GST-BRD4 BD1 Query titration to potential nucleosome Targets; dashed line represents EC_80_^rel^ (5 nM) against ([H4K5acK8acK12acK16ac]) (*aka*. ([H4tetra^ac^])). *[Main]* GST-BRD4 BD1 (5 nM) discovery screen to indicated panel reveals strong preference for ([H4tetra^ac^]) nucleosomes. **(D)** *[Inset]* GST-BRM BD titrations to potential nucleosome Targets; dashed line represents EC_20_^rel^ (30 nM) against ([H3K4acK9acK14acK18ac]) (*aka*. ([H3tetra^ac^])). The binding at near µM concentrations to unmodified nucleosomes is likely due to interactions with nucleosomal DNA [60] (**Suppl. Fig. 4C**). *[Main]* GST-BRM BD (30 nM) discovery screen to indicated panel identifies strong preference for ([H3tetra^ac^]) nucleosomes. See **Suppl. Figs 2-4** for Captify-Alpha two-dimensional (2D) titrations with a range of lysine-acyl readers, and **Suppl. Table 2** for complete datasets from peptide (n = 287) and nucleosome (n = 77) target discovery screens.

Of particular utility, Captify-Alpha is also compatible with defined nucleosome Targets, where continued improvements in synthesis and scaling have dramatically increased the diversity now available (24,26,66). In this version of the workflow, a regular first step is Query titration to potential negative and positive controls (usually unmodified and predicted engaged PTM(s)) (28), which can include an exploration of buffer components (*e.g.,* salt) and supplements (*e.g.,* exogenous DNA, cofactors, or divalent cations) to optimize S/B. Pinpointing a Query concentration between EC_20_^rel^ and EC_80_^rel^ with robust S/B is considered ideal for further use, per HTS guidelines (67). As an example, we tested GST-tagged forms of two disease relevant bromodomains that respectively engage acetylated H4 and H3 histone tails: BRD4 BD1 (31,65) and BRM BD (68–70). In each case, titration identified optimal reader concentrations (**Figs. 1C-D**, *Inset:* S/B ≥ 10) for discovery screening to nucleosomes (77-member panel; **Suppl. Table 2**), where each bromodomain preferred multi-acetylated Targets (**Figs 1C-D**, *Main*). Indeed, BRD4 BD1 failed to engage single H4 tail acetylations in the nucleosome context despite robust binding to comparable peptides (compare **Figs 1C** and **1B**), while BRM BD only weakly bound ([H3K14ac]), contrasting with prior peptide studies (69).

The restriction of reader : PTM binding preference on nucleosomes *vs*. peptides is commonly reported (27,28). To investigate further, we used Captify-Alpha to examine six additional acyl-readers, including two more bromodomains (BRD3 BD1 and BRG1 BD) and all four human YEATS domains: a family with strong links to transcriptional regulation and disease (71–73). For each Query, we established reader capability followed by a discovery screen with PTM-defined peptides; then optimized binding conditions to nucleosomes followed by a discovery screen to this target class. In every case, we observed refined binding on nucleosomes *vs*. peptides. Further, each reader preferred distinct conditions to yield an acceptable S/B (**Suppl. Figs 2 – 5**), which may inform on their individual means of Target engagement, but also stresses the importance of extensive optimization in such studies.

### Nucleosome context refines the binding preference of methyl-lysine readers

Having established assays for readers of lysine acylation, we extended our investigation to those of histone lysine methylation. This PTM class potentially exists at target residues in three major forms (me1, me2 and me3), and is regulated by families of writer and eraser enzymes (lysine methyltransferases and demethylases: KMTs and KDMs), with the distinct product states distinguished by genomic distribution and function (74–78).

Polycomb Group (PcG) protein L3MBTL1 contains three MBT (malignant brain tumor) repeats with a reported preference for lower lysine methylations (Kme1 or Kme2) on multiple linker and core histone residues, including H1.5K27, H3K4, H3K9, H3K27 and H4K20 (79–84). Query titration of GST-L3MBTL1 MBT to H4_[11-27]_K20 peptides (me0-1-2-3) confirmed preferential reader binding to the mono and dimethyl states (**Fig. 2A**), with discovery testing (at EC_80_^rel^; 3 nM) to the lysine-methyl status panel (K-MetStat; me0-1-2-3 at H3K4, H3K9, H3K27, H3K36, and H4K20) confirming interaction to each Kme1/Kme2 peptide (**Fig. 2B**). Such PTM state preference but residue promiscuity is due to a loose binding mode, where the MBT domain extensively interacts with the mono- or dimethyl-lysine without contacting surrounding residues (82,83). However, it would seem of questionable biological utility since each of these PTMs is distinctly regulated *in vivo* (74–78). We thus explored L3MBTL1 MBT binding to the K-MetStat nucleosome panel, and observed no engagement (**Fig. 2C**). This may suggest either that additional elements are required for L3MBTL1 MBT binding in the chromatin context, or the peptide data is of limited *in vivo* significance. Given the cancer relevance of L3MBTL1 (85,86), these findings warrant further investigation.

**Fig. 2.**
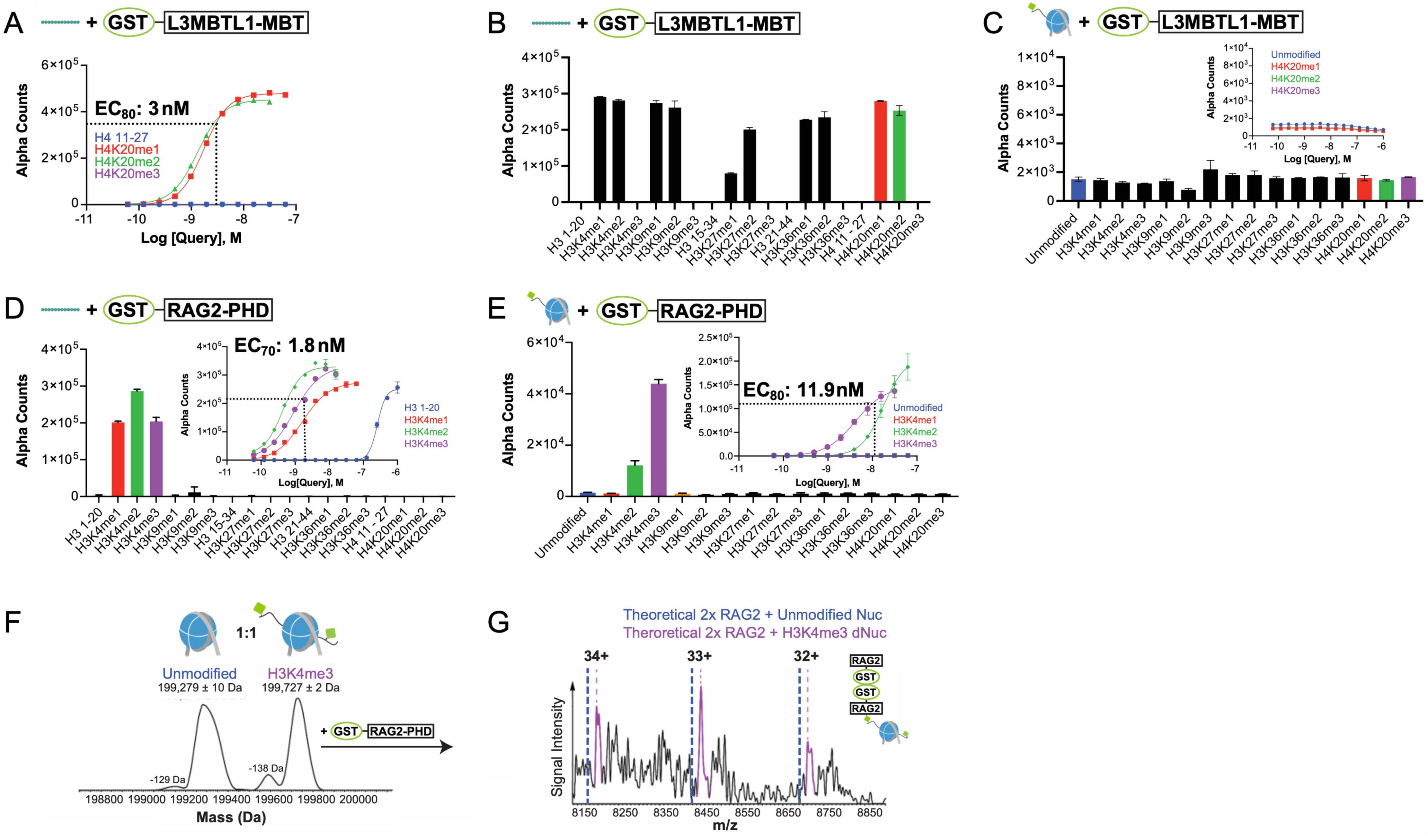
Lysine-methyl readers show refined PTM specificity in nucleosome *vs.* histone peptide assays. **(A)** GST-L3MBTL1 MBT Query titration to H4_[11-27]_K20 methyl peptide Targets (maximal S/B at EC_80_^rel^ = 3 nM) identifies the reported preference for Kme1 and Kme2. **(B)** GST-L3MBTL1 MBT (3 nM) discovery screen against 287-member peptide panel identifies a preference for Kme1 and Kme2 independent of histone residue (data subset shown). **(C)** *[Inset]* GST-L3MBTL1 MBT titration to nucleosome Targets bearing distinct methyl states at H4K20 identifies no binding preference. *[Main]* GST-L3MBTL1 MBT (3 nM) discovery screen shows background binding to nucleosomes independent of lysine-methyl status. **(D)** *[Inset]* GST-RAG2 PHD titration to H3_[1-20]_K4 methyl peptides (EC_70_^rel^ = 1.8 nM). *[Main]* GST-RAG2 PHD (1.8 nM) peptide discovery screen shows selective binding to methylations at H3K4 (me1-2-3) over all other histone methyl states (data subset shown). **(E)** *[Inset]* GST-RAG2 PHD titration to nucleosomes with H3K4 methyl states (EC_80_^rel^ = 11.9 nM). *[Main]* GST-RAG2 PHD (11.9 nM) nucleosome discovery screen reveals a refined preference for ([H3K4me3]). For **(B-E)**, see **Suppl. Table 2** for complete discovery screen datasets. **(F)** Native top-down mass spectrometry (nTDMS) distinguishes a 1:1 mix of unmodified and ([H3K4me3]) nucleosomes (both 1 μM). **(G)** GST-RAG2 PHD (5 μM) selectively associates with ([H3K4me3]) over unmodified nucleosomes. Reader was incubated with a 1:1 nucleosome mix (both 1 μM) and analyzed by nTDMS. Mass-to-Charge (m/z) peaks correspond to dimerized GST-RAG2 PHD bound to ([H3K4me3]) nucleosomes (further characterized in **Suppl Fig. 6**).

RAG complex (Recombination Activating Gene: RAG1-RAG2) is critical for adaptive immunity (87–89) and mediates DNA cleavage during V(D)J recombination at immunoglobulin and T-cell receptor genes. Within RAG, the plant homeodomain (PHD) of RAG2 binds H3K4me2/me3 for effective genomic targeting (90–92). To explore any impact of target context, we titrated GST-RAG2 PHD Query to H3K4methyl-focused peptides and nucleosomes (**Fig. 2D-E**, insets), followed by discovery screens to the respective K-MetStat panel. In peptide testing the reader bound all three methyl states (me1-2-3) over the unmodified residue at H3_[1-20]_K4 (**Fig. 2D**); while nucleosomes refined binding to a preference for the higher methylations (H3K4me3 > me2: **Fig. 2E**). Orthogonal validation of this engagement was provided by mixing unmodified and ([H3K4me3]) nucleosomes (1:1), confirming their distinct identities by native top-down mass spectrometry (nTDMS) (93,94) (**Fig. 2F**), then adding GST-RAG2 PHD and confirming the specific formation of a (GST-RAG2 PHD: ([H3K4me3])) complex (**Fig. 2G** and **Suppl. Fig. 6**).

We continued our analysis of methyl-lysine readers, selecting Queries to represent a diversity of domain types, targets and functions: chromodomains (CDs) from heterochromatin protein HP1β (CBX1) (95–97) and Polycomb protein CBX7 (98–102); PWWP domains from histone KMTs NSD2 and NSD3 (36,37), DNA methyltransferase DNMT3A (60,103), and non-enzymatic nuclear factors GLYR1 (104) and BRD1 (105); and the ADD domain of ATRX, a SWI/SNF chromatin remodeler (106,107). In each case Query engagement was established / optimized on PTM-defined peptides and nucleosomes, followed by discovery screens where each reader showed refined binding on nucleosome Targets. For example, GST-HP1β CD and GST-ATRX ADD engaged H3_[1-20]_K9 peptides containing all three methyl states (me3/2>1), but only ([H3K9me3]) nucleosomes (28) (**Fig. 3A-B** and **Suppl. Figs 7-9**). As expected, the PWWP domains largely failed to bind histone peptides (28,37), with the notable exception of DNMT3A which preferred H3_[21-44]_K36me2/3 but also engaged H3_[15-34]_K27me2/3 (**Suppl. Fig. 7A**). However, on nucleosomes all PWWPs showed varying selectivity for the higher methyl states of H3K36, which in many cases (*e.g.*, GLYR1: see below) free DNA competitor was required to discern (28) (**Fig. 3C-E** and **Suppl. Figs 7-9**).

**Fig. 3.**
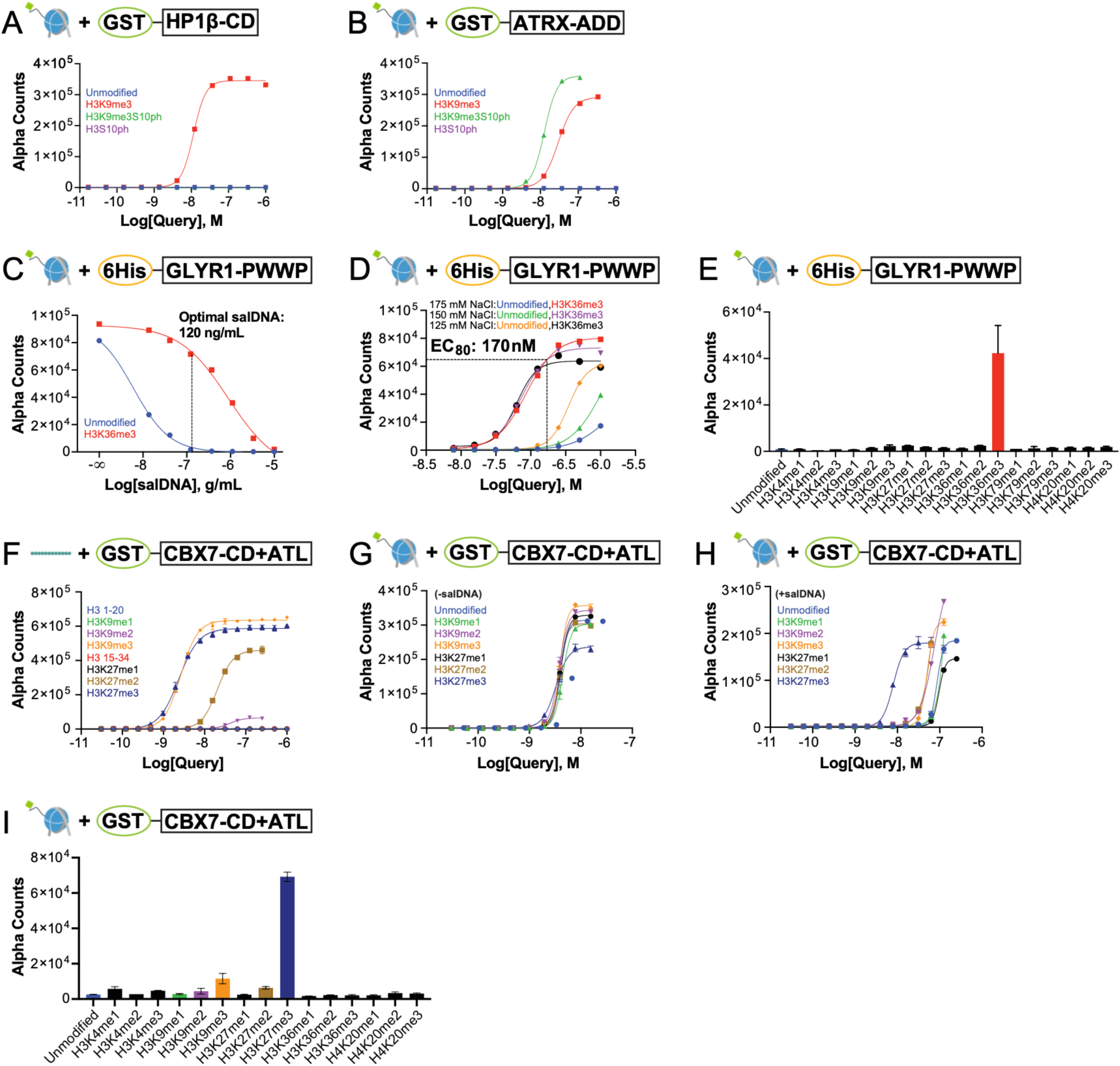
Nucleosome context is required for an accurate determination of multivalent reader engagement. **(A-B)** Titration of GST-HP1β (CBX1) CD **(A)** and GST-ATRX ADD **(B)** to indicated nucleosomes identifies differential impact of the ([H3K9me3S10ph]) combinatorial relative to ([H3K9me3]). **(C)** Salmon sperm DNA (salDNA) optimization for 6His-GLYR1 PWWP. Nonspecific nucleosome binding is reduced by the free DNA, revealing a preference for ([H3K36me3]). Dashed line represents optimal salDNA concentration (120 ng/mL) used for **(D-E)**. **(D)** Salt optimization for 6His-GLYR1 PWWP. Query was titrated at noted NaCl concentrations against ([H3K36me3]) and unmodified nucleosomes, where increasing salt reduced binding to latter (EC_80_^rel^ = 170 nM). **(E)** GLYR1 PWWP (170 nM) binding in a nucleosome discovery screen with optimized buffer conditions (120 ng/mL salDNA, 175 mM NaCl), confirms a preference for ([H3K36me3]). **(F)** GST-CBX7 CD-ATL titration to histone peptides identifies equivalent binding to H3_[1-20]_K9me3 and H3_[15-34]_K27me3. **(G-H)** CBX7 CD-ATL titration to nucleosomes reveals a preference for ([H3K27me3]), but only in presence of salDNA competitor (123 ng/mL: compare **G** and **H**) (EC_50_^rel^ = 7.1 nM). **(I)** CBX7 CD-ATL (7.1 nM) binding in a nucleosome discovery screen with optimized buffer conditions (120 ng/mL salDNA), confirms a preference for ([H3K27me3]). See **Suppl. Figs 7-9** for Captify-Alpha 2D optimizations of a range of lysine-methyl readers; **Suppl. Figs 11-12** for further dissection of the CBX7 CD-ATL tandem; and **Suppl. Table 2** for complete datasets from all discovery screens.

### The unpredictability of methyllysine analogs (MLAs) for binding studies

Structural analogs of the Kme1-3 states (K_C_me1-3; methyl-amino-alkylated cysteines (108–110) (*e.g.*, **Suppl. Fig. 10A**)) continue to be employed for binding and structural studies, most particularly in NMR spectroscopy, which regularly uses ^15^N-labeled recombinant histones for K_C_me conversion. Despite this technical convenience, the value of such studies is undermined if the MLA does not functionally behave as the structurally similar native Kme. To investigate, we created nucleosomes with native H3K4me3 and H3K9me3 and their MLAs (respectively H3K4_C_me3 and H3K9_C_me3), and compared antibody and reader binding to each class. Here an anti-H3K4me3 and anti-H3K9me3 bound the native methyl and MLA with similar efficiencies, while the readers showed contrasting tolerances: RAG2 PHD bound equivalently to ([H3K4me3]) and ([H3K4_C_me3]), while ATRX ADD effectively engaged ([H3K9me3]) but was blind to ([H3K9_C_me3]) (**Suppl. Fig. 10C-F**). Such observations are a salutary reminder of the unpredictable nature of MLAs, which often support weaker reader domain engagement relative to native methylations (111–114), and where the safest path appears to directly compare reader binding with each entity before adopting MLAs for technical convenience.

### Interrogating the impact of histone PTM crosstalk and multivalent engagement

Chromatin functionality through PTMs requires coordinated engagements, where various elements can have a positive or negative impact (14,22,24,48,115). As an example, the HP1β CD and ATRX ADD reader domains both interact with H3K9me3 to regulate gene silencing and genome stability (106,107,116–119). However, their binding is differentially impacted by H3S10 phosphorylation in *cis*, which displaces HP1β CD (116,117) while ATRX ADD is unaffected (118,119). This mechanism was recapitulated by Captify-Alpha with nucleosome Targets, where combinatorial S10ph abolished HP1β CD binding to H3K9me3, but three-fold enhanced that of ATRX-ADD (**Fig. 3A-B**).

CAP complex : nucleosome engagement is largely (if not invariably) multivalent, with multiple contact points on the histone tails (*e.g.*, at PTMs in *cis* or *trans*) and other nucleosome surfaces, such as the H2A/H2B acidic patch and/or DNA (by charge, sequence motif or modification). Such modes of engagement are exhibited by individual reader domains with multimodal binding ability (*e.g.,* the PTM and DNA binding PWWPs (57)), or grouped reader domains that independently bind co-occurring PTMs in *cis* and/or *trans*, such as the BPTF PHD-BD (27,47) or PHIP/BRWD2 Tudor-BD1-BD2 (59).

PWWP domains have a general preference for H3K36me2/me3 (**Suppl. Fig. 7**), but many also bind DNA (120). As an example, we examined the PWWP domain from GLYR1, a cofactor of the H3K4me3 demethylase LSD2 (121,122) and H3K36me3 reader (104) with putative roles in transcriptional elongation (123,124). In initial testing 6His-GLYR1 PWWP showed equivalent binding to unmodified and ([H3K36me3]) nucleosomes, but a profound preference for the latter in the presence of salmon sperm DNA (salDNA) competitor (**Fig. 3C**), that was further improved by salt optimization (**Fig. 3D** and **Suppl. Fig. 9G**). Screening the K-MetStat nucleosome panel with these optimized conditions confirmed H3K36me3 as the primary Target (**Fig. 3E** and **Suppl. Fig. 7B**).

Polycomb Repressive Complex 1 (PRC1) co-operates with PRC2 to establish repressive H3K27me3 heterochromatin domains that regulate gene expression, development and higher-order chromatin architecture (125). The PRC1 CBX7 subunit contains a H3K27me3-binding chromodomain (CD) (101,126–128) and adjacent DNA-binding AT-hook-like (ATL) motif (98–102). Both are required for stable chromatin interaction *in vivo* and proposed to act as a functional unit, with initial H3K27me3 engagement of the chromodomain triggering an allosteric change that enables the ATL to engage DNA (101). However, CBX7 CD is also reported to bind equivalently to H3K9me3 and H3K27me3 (98–100), a capability of unknown relevance.

Captify-Alpha was used to examine Target binding by diverse GST-CBX7 queries, including the individual CD or ATL, the tandem CD-ATL, and a tandem containing a loss-of-function point-mutated CD (101) (CBX7 CD(F11A)-ATL). Studies with histone peptides (H3_[1-20]_ or H3_[15-34]_), revealed equivalent binding of CBX7 CD to K27me3 and K9me3 > H3K27me2 ~ H3K9me2 (**Suppl. Fig. 12A**), while CBX7 CD-ATL bound H3K27me3 ~ H3K9me3 > H3K27me2 >> H3K9me2 (**Fig. 3F**), and the chromodomain mutation ablated all binding (**Suppl. Fig. 12D**). So, the CBX7 CD mediates effective and indistinguishable binding to trimethylated H3K9 and H3K27 in the peptide context, possibly driven by the sequence similarity of these regions (…AR[K9]S<u>T</u>… *vs* …AR[K27]S<u>A</u>…): a distinction difficulty shared by many antibodies ostensibly to the methyl states of K9 or K27 (chromatinantibodies.com/). On peptides the CBX7 ATL is of minor consequence, somehow contributing to selectivity between the dimethyl states.

We next investigated the binding of CBX7 CD to PTM-defined nucleosomes. Here the higher order context refined chromodomain binding preference to ([H3K27me3]) over other methyl states at H3K27 and H3K9 (**Suppl. Fig. 12B**), consistent with the CBX7 role in polycomb-repressed chromatin. The native context for CBX7 CD is with the adjacent DNA-binding ATL, but the tandem constructs (CD-ATL or CD(F11A)-ATL) were dominated by DNA binding, with no selectivity between unmodified and methylated nucleosomes (**Fig. 3G** and **Suppl. Fig. 12E**). However, salDNA competitor revealed the preference of CD-ATL for ([H3K27me3]) (**Fig. 3H** and **Suppl. Figs 11 – 12**), that was further demonstrated on testing the tandem reader to K-MetStat nucleosome panel under optimized conditions (EC_50_^Rel^: **Fig. 3I**). In this manner, the CBX7 CD-ATL tandem uses a multivalent mechanism to engage nucleosomes, with the higher order context also conferring a chromodomain specificity not displayed on histone peptides. This is in general agreement with a model where the CBX7 CD-ATL comprise a functional unit to engage chromatin (101), and reconciles conflicting published data, where all suggesting CBX7 CD bound to H3K9me3 used histone peptides.

### Development of a wash-based multiplex platform that predicts genomic mapping capability for reader binding studies

No-wash Captify-Alpha can effectively profile CAP : nucleosome binding preferences. However, the approach also requires adaptations to interrogate multivalent binding, such as inclusion of salDNA to isolate a DNA-mediated element (*e.g.*, **Figs. 3C** and **3H**). Stringency can also be increased by washing, spurring our development of Captify^TM^-Luminex (**Fig. 4A**). This offers numerous advantages: high sensitivity, reproducibility and throughput, multiplex capabilities (*i.e.,* testing multiple Targets in a single Query reaction), and compatibility with CUT&RUN wash conditions, offering a cost-effective means to optimize reader binding before genomic analyses.

**Fig. 4.**
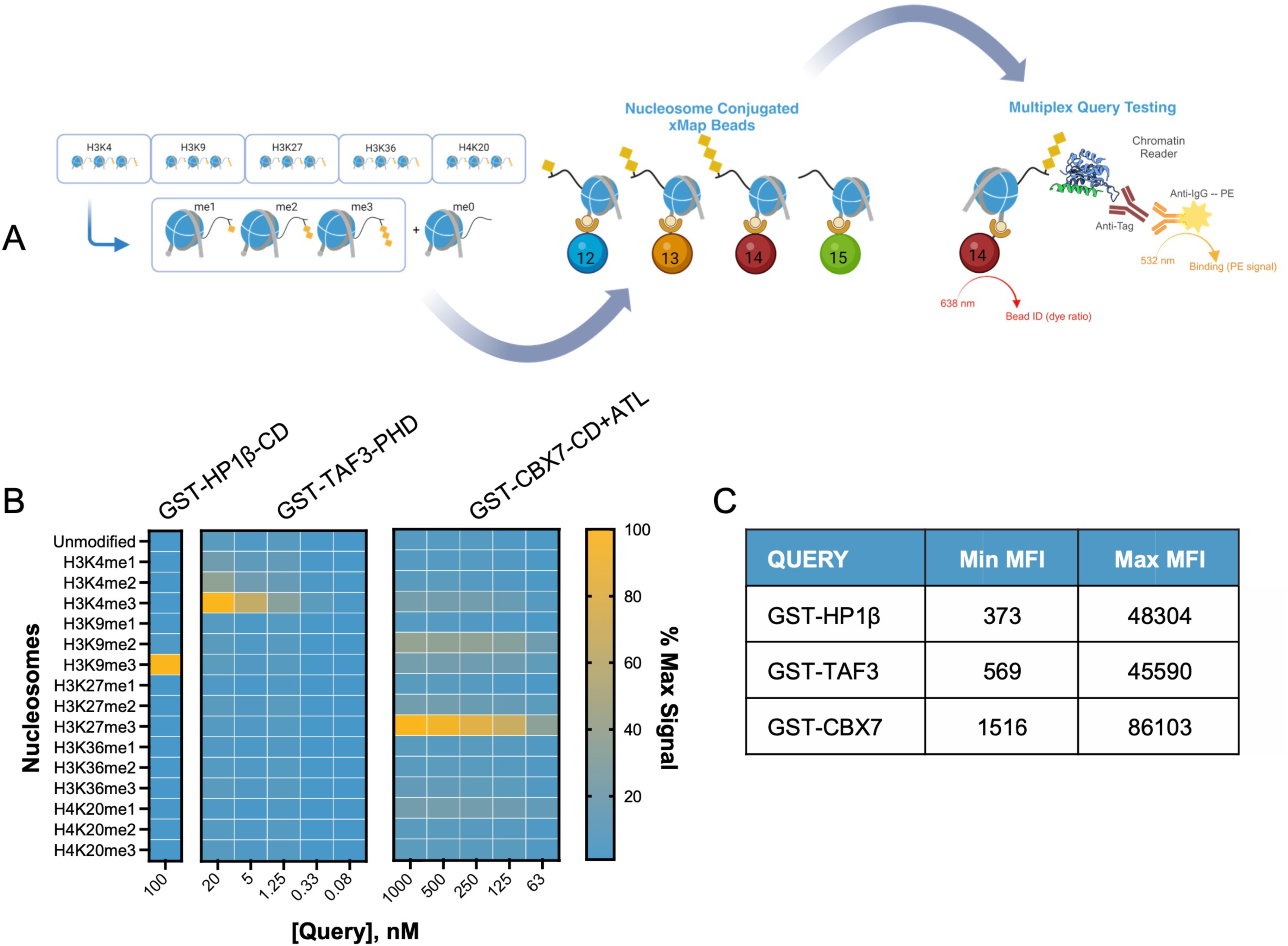
Captify-Luminex enables interrogation of chromatin reader binding to multiplexed nucleosome Targets. **(A)** Biotinylated nucleosome Targets (the K-MetStat panel: me0-1-2-3 at H3K4, H3K9, H3K27, H3K36 and H4K20) are individually conjugated to distinct Luminex avidin-coated MagPlex bead regions, pooled, and probed in multiplex with GST-tagged reader domains. Interactions are detected using anti-GST and anti-IgG*PE secondary, with bead/PE signals measured on a FLEXMAP-3D System. **(B)** GST-HP1β CD, GST-TAF3 PHD and GST-CBX7 CD-ATL binding across the K-MetStat panel. For each Query concentration (nM; labeled columns), heatmap depicts signal as percentage of max median fluorescence intensity (MFI). **(C)** Signal range for each Query in **(B)**.

Luminex xMAP uses fluorescently barcoded MagPlex**^®^** microspheres (*aka*. bead regions) to bind distinct Targets and enable multiplexing. For Captify-Luminex, biotinylated PTM-defined nucleosomes (Targets) were individually conjugated to discrete MagPlex-Avidin bead regions (providing a fluorescent barcode of histone PTM identity), and then pooled to multiplex panels (*e.g.,* K-MetStat: me0-1-2-3 at H3K4, H3K9, H3K27, H3K36 and H4K20). GST-tagged Queries were then combined with the panel in 96-well plates, detected with a fluorescent label (using anti-GST and anti-IgG*phycoerythrin (PE)), and relative binding assessed by co-localization of Query (fluorescent PE) and Target (fluorescent bead region) (**Methods** and **Fig. 4A**). Notably, multiplexing also provides a competitive environment, reflective of studies with native chromatin.

As an initial capability test, we focused on an integral nucleosome surface: the acidic patch. This cluster of negatively charged residues in histones H2A and H2B is a central hub for nucleosome interactors, including diverse chromatin modifying and remodeling enzymes to regulate their activity, and the H4 tail to promote chromatin fiber formation (129–132). H2BK120ub partially occludes the acid patch, with this ‘gatekeeping’ function of the labile and mobile PTM a potential means to regulate access for pathway-specific patch binders within actively transcribed genes (63,133–136). Captify-Alpha and -Luminex were used to examine acid-patch binding by the VHH Chromatibody (137), and each approach confirmed effective engagement with unmodified nucleosomes but not an acid-patch mutant (H2AE92K) (**Suppl. Fig. 13A-B**). However, the ([H2BK120ub]) nucleosome was indistinguishable from unmodified in Alpha, but closer to the acid-patch mutant in Luminex (63) (**Suppl. Fig. 13A-B**). In this manner, acid-patch binding by chromatibody was stable enough to tolerate H2BK120ub in the no-wash platform, but PTM impact was revealed by the multi-step washable approach.

We next applied Captify-Luminex to three GST-tagged lysine-methyl readers : HP1β CD with a preference for H3K9me3 (**Fig. 3A**), TAF3 PHD with a preference for H3K4me3 (138,139), and CBX7 CD-ATL with a preference for H3K27me3 in the presence of salDNA (**Fig. 3H**-**I**). Each preference was recapitulated with the Luminex-approach, noting that CBX7 CD-ATL no longer required salDNA competitor to selectively engage H3K27me3. Additionally, each S/B was regulatable by Query concentration, and under optimal conditions provided a >55-fold ratio from target to unmodified nucleosome (**Fig. 4B-C**).

Previously, we established the reader-CUT&RUN method using the BPTF PHD-BD tandem, observing a high degree of overlap with its combinatorial PTM-targets ({H3K4me4K14acK18ac}) and endogenous BPTF (27). Supported by fully-defined DNA-barcoded spike-in nucleosome controls, reader-CUT&RUN enables the *in vivo* dissection of binding preference for chromatin features, including combinatorial histone PTMs, DNA modifications, and higher-order structures. Building on this, we considered that Captify-Luminex optimized conditions might be further suitable for the genomics approach. In this manner each GST-tagged lysine-methyl reader manifested spike-in nucleosome recoveries and genomic enrichment patterns similar to an antibody to their ostensible PTM targets: TAF3 PHD / anti-H3K4me3 formed sharp peaks at active promoters; CBX7 CD-ATL / anti-H3K27me3 spanned polycomb-repressed genes; and HP1β CD / anti-H3K9me3 covered broad regions across constitutively repressed locations (**Fig. 5** and **Suppl. Fig 15**).

**Fig 5.**
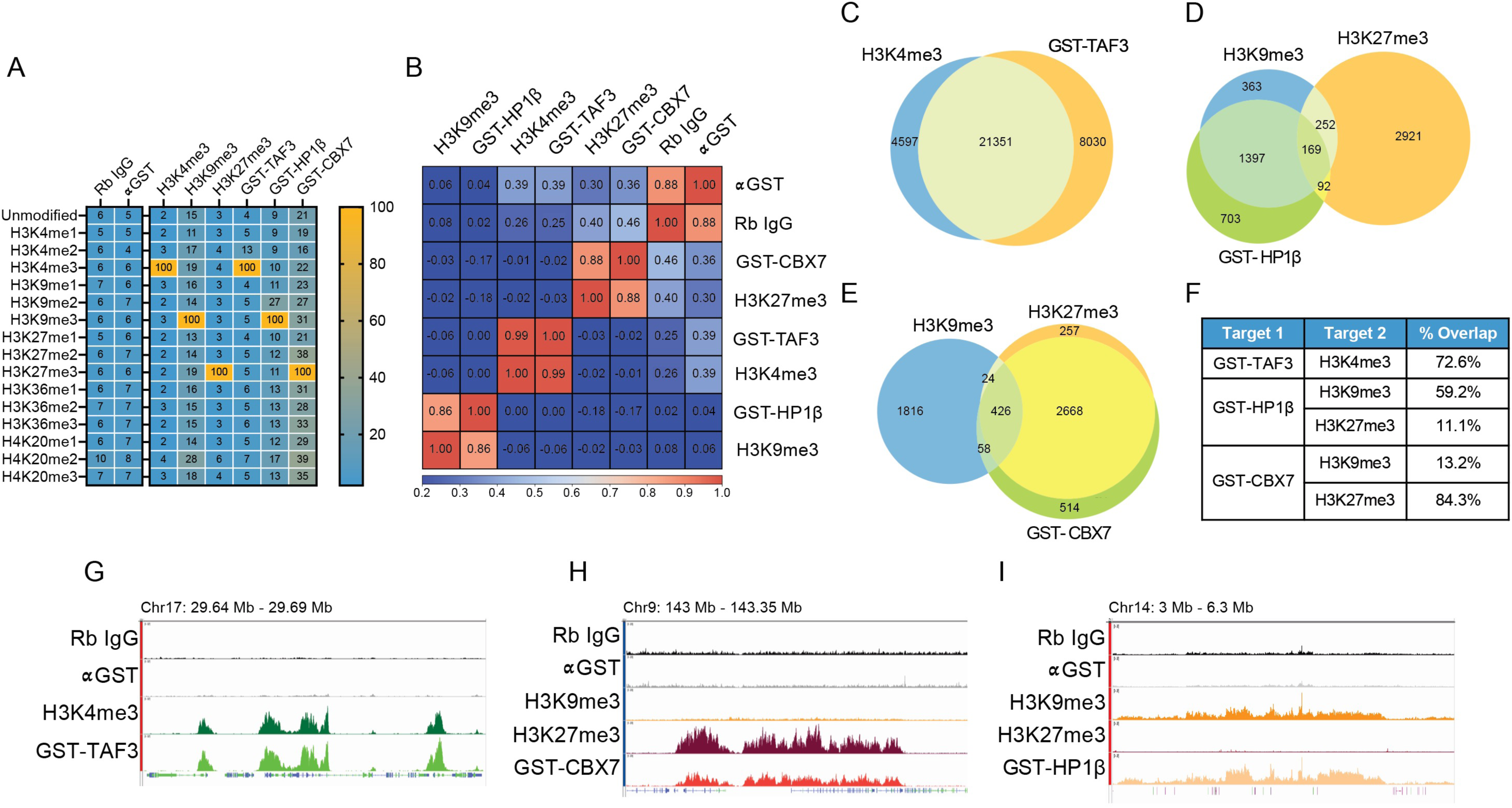
Readers can deliver antibody-like PTM profiling in CUT&RUN genomic mapping. **(A)** DNA-barcoded nucleosome spike-in (K-MetStat panel) recovery from Reader-CUT&RUN reactions. Columns group data by antibody / GST-tagged reader; rows depict recovery of each PTM in K-MetStat panel normalized to predicted Target (100%; orange in heatmap), except for anti-IgG and anti-GST (% of total spike-in reads). **(B)** Pearson correlation plot compares CUT&RUN read overlap from antibodies *vs.* GST-Readers (1 = perfect correlation). **(C-F)** Venn overlap of called peaks between PTM-specific antibodies and GST-Readers (≥ 50% peak overlap classed as positive). Table shows average peak width and % overlapping peaks in each pairwise comparison (Target 1, GST-Reader; Target 2, PTM-specific antibody). **(G-I)** Representative peak comparisons: H3K4me3 and GST-TAF3 PHD **(G)**; H3K9me3, H3K27me3 and GST-CBX7 CD-ATL **(H)**; H3K9me3, H3K27me3 and GST-HP1β CD **(I)**. Each IGV browser window (genomic region noted) is group scaled to * track. IgG and αGST are background controls.

## DISCUSSION

Accurate interrogations of chromatin reader interactions are crucial to advancing our understanding of normal biology and the disease state. In this work we develop, validate and integrate a range of biochemical and genomic mapping approaches to study chromatin readers in the nucleosome context. Direct comparison showed histone peptides often failed to accurately predict the nucleosome binding preferences of even single reader domains, and were uninformative or even misleading for multivalent engagement, particularly when a DNA binding component was involved (*e.g.*, **Figs 1-3**). In stark contrast, *in vitro* performance in Captify-Luminex could be applied to CUT&RUN genomic mapping, where various lysine-methyl readers displayed antibody-like capability to distinguish various histone lysine-methyl states (*e.g.*, **Figs 4-5**). This offers a path to the more efficient study of novel chromatin regulators and their disease-associated mutations.

### The limitations of surrogates

Histone PTM peptides have been truly enabling for decades of chromatin studies (24). Their primary strength lay in convenience: fully PTM-defined histone regions at an accessible price, control over experimental variables, and the ability to consistently generate data. All this in the context that PTM-defined nucleosomes were often unavailable or largely inaccessible. However histones peptides lack the structural complexity required to fully explore CAP interactions (35–46), and any results can even be directly misleading (28,55) (*e.g.*, **Figs. 1-3**). Further, fully defined nucleosomes are now readily available and fully compatible with biochemical, proteomic, structural and genomic approaches (26,28,55,59–63,66,94,111,129,140–144). Here we perform extended analyses on multiple CAP domains that engage PTMs, DNA or the acidic patch. Some general observations emerge, including: that nucleosomes consistently refine binding preferences relative to histones peptides; that multivalent nucleosome engagement by CAPs is more often regulatory than additive; and that while reductive assays with surrogates (*i.e.*, isolated domains, histones peptides, or MLAs) can yield data, it too often conflicts with data from multi-domain CAPs, nucleosomes, and native PTMs. Given this, the most valid mechanistic insight most likely resides in the more complicated systems and a move away from the surrogates is largely warranted.

### New insights await, but any exploration of nucleosome engagement must accommodate the multivalent milieu

Reader domain : nucleosome binding studies often require extensive optimization (*e.g.*, the 2D-Query concentration and salt explorations of **Suppl. Figs 4** and **9**, or competitor DNA titrations of **Fig. 3C,G,H** and **Suppl. Figs 11** – **12**). Here a major strength of no-wash Captify-Alpha is its flexibility for modified conditions to support weak binders removed from their native context (*e.g.*, **Suppl. Fig. 4**), or to isolate a particularly dominant element (*e.g.*, the DNA binding AT-like hook motif in the CBX7 CD-ATL tandem: **Fig. 3F-I** and **Suppl. Figs 11** – **12**); though it proved unable to reveal H2BK120ub-attenuated binding of chromatibody VHH to the acidic-patch (**Suppl. Fig. 13**). However, a major strength of washable and competitive Captify-Luminex is using its higher stringency to explore multivalent engagement. As an example, the approach had no difficulty identifying the H3K27me3 preference of CBX7 CD-ATL without competitor DNA (**Fig. 4**), and even suggested optimized conditions where the recombinant reader was capable of mapping H3K27me3 with similar metrics to a PTM-specific antibody in CUT&RUN (**Fig. 5** and **Suppl. Fig 15**).

Our analyses of individual bromodomains often revealed a strong preference for multiply modified nucleosome substrates, despite each BD generally only being able to accommodate a single acetylated lysine in their PTM binding pocket (31,145–148). For example, BRD4 BD1 displayed strong binding to ([H4K5acK8acK12acK16ac]) but none of the single H4 acetylations (**Fig. 1C**), while BRM BD had >ten-fold stronger binding to ([H3K4acK9acK14acK18ac]) over ([H3K14ac]) (**Fig. 1D** and **Suppl. Table 2**). This could represent increased target accessibility, where multiple acetylations weaken the histone tail interaction with nucleosomal DNA (28,55,132,149–154), but only one acetylated residue is primarily bound by the BD. Alternatively high local concentrations of acetylated lysines (as frequently observed on *in vivo* chromatin (147,155)) could contribute transient engagements to the recruitment and/or stabilization of a site-specific Kac binding event. However, individual bromodomains are almost certainly not biologically expected to manifest an individual role. As an example, BRD4 contains two adjacent bromodomains (BD1 and BD2), with the reader tandem effectively enriching native nucleoforms containing di- and tri-acetylated histone H4 tails (156). Further, while BPTF BD weakly binds ([H3K4acK9acK14acK18ac]), the PHD-BD tandem binds similarly to ([H3K4me3K9acK14acK18ac]) and ([H3K4me3K14ac]) (27). In each case combinatorial context synergizes at least two individually weak site-specific interactions, likely representing the biological mechanism of action. With this in mind we used the GST-epitope tag through this study to take advantage of its dimerization (157), and thus provide an ‘avidity boost’ for target engagement, just as prior work did by multimerizing the HP1β CD (158). However, while useful for nucleosome studies, this is not a necessity and non-dimerizing epitope tags (*e.g.*, MBP, 6HIS, FLAG or HA) have also been successfully employed (28,59–61,158).

### So, can we study everything now?

Combinatorial interactions are inherent to many CAPs, enabling robust and specific binding to their chromatin targets. These interactions typically involve two or more CAP moieties engaging distinct PTMs, histone sequence motif, DNA (by general charge, position on the nucleosome surface, sequence motif or modification), nucleosome surfaces (*e.g.*, the H2A/H2B acidic patch), or indeed other CAPs. Evaluating such complex interactions offers a formidable challenge as the study platform(s) must be sufficiently sensitive and quantitative to isolate and evaluate individual contributions. Here we describe easily adoptable biochemical and genomic platforms with such capabilities. Further, the tools now exist to create highly defined nucleoforms, including those containing histone mutants, PTMs or combinatorials, in *cis* or *trans*, alongside DNA sequence motifs or modifications, and is stepping towards synthetic chromatin arrays that even more closely represent physiological targets (56,159–161): all in the service of dissecting biological mechanism with ever greater precision.

The new approaches will eventually hit a limit when interrogating multi-subunit CAP complexes and chromatin arrays: how many points of *cis* / *trans* contact can they reveal? There appears a lot of space to explore: Captify-Alpha can discern all partners in a trivalent interaction between PHIP/BRWD2 Tudor-BD1-BD2 and ([H3K4me3K14ac]•[H4K12ac]) nucleosomes, and indeed showed the impact of disease-associated mutations on chromatin-binding [54]. The use of reader domains for genomics is particularly exciting, since tandems can find combinatorial PTM signatures; either native (*e.g.*, BPTF PHD-BD as above) or by design (*e.g.*, the DNMT3A-MPP8 PWWP-CD chimera that enriches ({H3K9me3K36me3})) (156,162,163) that overlaid signals from PTM-specific antibodies can only infer. As such, the current technologies offer a highly accommodating starting point for future discoveries.

## **SUPPLEMENTARY MATERIAL** associated with this manuscript

2x Supplementary tables (each with multiple tabs)

15x Supplementary Figures

Supplementary References

## MATERIALS & METHODS

### Histone peptides

PTM-defined peptides for Captify™ assays were synthesized with a terminal PEG-Biotin (locations as indicated: **Suppl. Table 1**) and identity / purity confirmed by mass spectrometry (MS).

### Semi-synthetic nucleosomes

Fully PTM-defined histones, octamers and nucleosomes (dNucs™ or versaNucs™: **Suppl. Table 1**) for Captify were synthesized, purified, and assembled on 147bp 5’ biotinylated 601 DNA (*EpiCypher* 18-0005) as previously (32), but without DNA barcoding.

For dNucs, PTM-defined histones were mixed to defined stoichiometry, dialyzed, purified to octamers, and assembled onto 147bp 5’ biotinylated 601 DNA [135]. The resulting products (*e.g.,* H3K27me3; *EpiCypher* 16-0317) contained full-length ‘scarless’ histones and minimal (<5%) free-DNA. PTMs were confirmed by MS and immunoblotting (if an antibody was available).

For versaNucs (27,55), histone H3 tail peptides (aa1-31; A29L) with a designation of interest (PTM, mutation or MLA) were individually ligated to a H3 tailless nucleosome precursor (*e.g.,* H3.1NΔ32; *EpiCypher* 16-0016). The resulting nucleosomes (assembled at 20-100 μg scale) contained undetectable levels of free peptide, and ≥90% full-length H3.1 with the designation of interest. When interrogating readers and CAPs in this study, we observed no difference between dNuc and versaNuc behavior and thus used these substrates interchangeably. Of note, versaNucs contain A29L and are not recommended for studies near this position (*e.g.,* modifiers / binders of H3R26, K27, or S28).

For nucleosomes with methyllysine analogs (MLA) histone H3 peptides (aa1-31; A29L) containing K4C or K9C were site-specifically reacted with the corresponding haloalkylamine: (2-bromoethyl) trimethyl ammonium bromide for K_C_me3; 2-chloro-N,N-dimethyl-ethylamine hydrochloride for K_C_me2; or 2-chloroethyl(methyl)ammonium chloride for K_C_me1; under SN2 reaction conditions (55,109,110) and purified for individual ligation to H3.1NΔ32 by versaNuc approach.

### Query proteins

Reader domains were recombinantly expressed as GST- or 6His-fusions (**Suppl. Table 1**), and purity assessed after SDS-PAGE and Coomassie staining (**Suppl. Figs 5** and **14**). Single domain VHH (Variable Heavy Domain of a Heavy chain antibody) Chromatibody to the nucleosome acidic patch was recombinantly expressed with a C-terminal 6His-tag and epitope characterized as previously (63,137).

### EpiTriton™ Histone Peptide Arrays

Three identical arrays of biotinylated histone peptides (287X (**Suppl. Table 1**) in two sets of triplicate plus a fluorescein transfer control) were printed on streptavidin-coated glass slides (EpiTriton™; *EpCypher* 11-4001) and stored / handled under subdued lighting. Prior to use, each slide subarray was submerged in hybridization buffer (1X PBS, 5% BSA, 0.1% (w/v) Tween-20), placed in a humidified chamber and incubated for 30 mins at 4 °C with gentle rotation. The liquid on each subarray was replaced with GST-tagged Query (2 μM in hybridization buffer) and incubated for three hrs at 4 °C with gentle rotation. Slides were washed three times with 1X PBS in a rotating slide chamber at 4 °C. To detect Query binding, slides were probed with primary anti-GST (*Sigma* G7781) for two hrs, washed, and secondary anti-rabbit IgG AlexaFluor 647 (*Invitrogen* A-21244) for 30 mins (all under subdued lighting in a humidified chamber at 4 °C with gentle rotation). After each step slides were washed with 1X PBS, and finally dried by centrifugation at 800x g for two mins. Fluorescent signal (526 nm and 670 nm) was measured on a Typhoon Trio+ (*GE*) using a +3 mm focal plane and 25 μm resolution. Signal was analyzed using ImageQuant TL (*Cytiva*) and Excel (*Microsoft*). All related data in **Suppl. Table 2**.

### Captify-Alpha

The assay previously known as dCypher™ is now named Captify™, with no distinction in how the assay is performed or its capabilities. Assays on the no-wash Alpha (Amplified luminescence proximity homogeneous assay) platform (*PerkinElmer*, *Revvity*) to examine the interaction of epitope-tagged reader domains (the Queries) with biotinylated PTM-defined peptides or nucleosomes (the Targets) were as previously (28,57,59,60). Importantly, Captify-Alpha assay buffers and bead binding buffers vary by Target type. When using peptides, assay buffer and bead binding buffer (50 mM Tris pH 7.5, 50 mM NaCl, 0.01% (v/v) Tween-20, 0.01% (w/v) BSA, 0.0004% Poly-L Lysine, 1 mM TCEP) were identical unless otherwise noted. In nucleosome-based experiments, assay buffer was (20 mM Tris (pH 7.5), 100-250 mM NaCl, 0.01% BSA, 0.01% NP-40, 1 mM DTT); bead binding buffer was the same minus DTT.

Queries and Targets were prepared in suitable assay buffer (as above). In a 384-well plate, 5 μL of GST- or 6His-tagged Query was combined with 5 μL of biotinylated peptide (100 nM final) or nucleosome (10 nM final) and incubated for 30 mins at room temperature. Anti-tag donor beads and streptavidin acceptor beads were prepared in appropriate bead binding buffers as above. For GST-tagged Queries, a 10 μL mixture of 2.5 μg/mL glutathione acceptor beads (*PerkinElmer* AL109) and 5 μg/mL streptavidin donor beads (*PerkinElmer* 6760002) was added to each well. For 6His-tagged proteins (including 6His-VHH to the nucleosome acidic-patch (63,137)), a 10 μL mix of 2.5 μg/mL Ni-NTA acceptor beads (*PerkinElmer* 6760619) and 10 μg/mL streptavidin donor beads was added. The plate was incubated at room temperature in subdued lighting for 60 mins, and Alpha signal measured on a *PerkinElmer 2104 EnVision* (680 nm laser excitation, 570 nm emission filter ± 50 nm bandwidth). Each reaction was performed in triplicate. Binding curves [Query : Target] were generated using a non-linear 4PL curve fit in GraphPad Prism 10, with EC_50_^rel^ values computed for specific comparisons. Where necessary, values beyond the hook point (indicating bead saturation / competition with unbound Query) were excluded and top signal constrained to average max signal for Target. In cases where signal never plateaued, signal was constrained to the average max signal within the assay. To optimize binding conditions for nucleosome assays, Query was cross-titrated in two-dimensions (2D) with salt (NaCl) or salmon sperm DNA (salDNA: *ThermoFisher Scientific* 15632011) against a predicted Target and unmodified nucleosome (<u>P</u>ositive and <u>N</u>egative (background) controls respectively). Optimal conditions produced robust signal-to-background (and were used for discovery studies within the range EC_20_^rel^ – EC_80_^rel^ as indicated), preferentially near physiological salt concentration in nuclei (~150 mM NaCl) [136]. Concentrations tested are indicated in respective figures (**Suppl. Figs 3, 4, 8** and **9**; see also **Suppl. Table 2**).

### Preparation of MagPlex nucleosome panels

Avidin-conjugated MagPlex^®^ beads (*Luminex, Diasorin*) with spectrally distinct regions were used to assemble multiplexed nucleosome panels (**Suppl. Table 1)**. The histone lysine methylation panel (K-MetStat) comprised 16 nucleosomes: me1, me2, and me3 on H3K4, H3K9, H3K27, H3K36, and H4K20 and unmodified control. The acidic patch assessment panel comprised six nucleosomes: H2AK119ub, H2BK120ub, H2AE61A, H2AE92K, H2AE105A/E113A, and unmodified control. All MagPlex bead handling and incubation was performed under subdued lighting. Briefly, beads stocks were vortexed and incubated for 30 sec in a water bath sonicator to monodisperse. Desired bead volumes were transferred to 1.5 mL tubes, placed on a magnet, and washed twice with pre-conjugation buffer (50 mM Tris pH 7.5, 0.01% Tween-20). Thoroughly mixed beads were added to the assigned nucleosome (*e.g.,* MagPlex Region 13 & ([H3K4me3])), adjusted to 500 μL (5 μg nucleosome per 1×10^6^ beads: *i.e.*, loading to saturation) with pre-conjugation buffer, and incubated on a rotator for 30 min at room temperature. Nucleosome-bound beads were washed twice with post-conjugation buffer (50 mM Tris pH 7.5, 0.01% Tween-20, 0.01% BSA), counted (LUNA cell counter; *Logos Biosystems*), and adjusted to 1×10^6^/mL. Beads were pooled to a master mix with an equal quantity (*e.g.,* 500,000 beads) of each MagPlex bead region, resuspended at 1×10^6^/mL per region in storage buffer (10 mM cacodylate pH 7.5, 0.01% BSA, 0.01% Tween-20, 1 mM EDTA, 10 mM beta mercaptoethanol, 50% glycerol), and stored at −20°C.

Panel balance, nucleosome integrity and [Nucleosome : bead region] identity were confirmed using anti-dsDNA (EMD Millipore #MAB030; 1/5, 1/50 and 1/500), anti-histone (*MilliporeSigma* MAB3422), anti-H3.1/2 (Active Motif #61629) and an appropriate anti-PTM if available (*e.g.*, <u>chromatinantibodies.com/</u>) (all **Suppl. Table 1**). Bead and antibody dilutions were prepared in QC assay buffer (50 mM Tris pH 7.5, 250 mM NaCl, 0.01% Tween-20, 0.01% BSA). Briefly, 50 µL of the multiplexed bead panel (20,000 beads/mL/region; 1,000 beads/region) was combined with 50 µL of antibody (1:125, 1:500 and 1:2000 unless otherwise specified) in a 96-well plate and incubated for 60 mins with shaking (800 rpm) to maintain bead resuspension. Beads were washed for three cycles on a magnet using 100 µL QC assay buffer, shaking for two mins between cycles. Anti-IgG PE (anti-rabbit, *Biolegend* 406421; or anti-mouse, *Biolegend* 405307) was diluted 1:100 in QC assay buffer, added to each well, and incubated for 30 mins with shaking. Beads were then washed for two cycles and resuspended in 100 µL QC assay buffer. Median fluorescence intensity (MFI) was measured using the FLEXMAP-3D System (*Diasorin*) with a minimum of 50 events/region. Bar graphs were generated in GraphPad Prism 10.

### Captify-Luminex

Assays to examine the interaction of epitope-tagged reader domains (the Queries) with biotinylated PTM-defined nucleosomes (the Targets) in the multiplex K-MetStat Luminex panel were performed in 96-well plates (GreinerBio 655900) with a modified CUT&RUN buffer (20 mM Hepes pH 7.5, 150 mM NaCl, 0.01% BSA, 0.01% Tween-20) or optimal Query buffer from Captify-Alpha (20 mM Tris pH 7.5, 150-250 mM NaCl, 0.01% BSA, 0.01% Tween-20, 1 mM DTT). Briefly, 50 μL multiplexed bead panel (20,000 beads/mL/region; 1,000 beads/region) was combined with 50 μL tagged Query (fixed concentration or titration; noted in figures / legends). The reaction plate was incubated for 60 min with shaking (800 rpm) to maintain bead resuspension. Beads were washed for three cycles on a magnet using 100 μL assay buffer, shaking for two min between cycles. 100 μL of anti-GST (1:2000; *Fortis* A190-122A) was added to each well and incubated for 30 mins with shaking. Beads were washed for three cycles and incubated with rabbit anti-IgG PE (1:100; *Biolegend* 406421) for 30 min with shaking. Beads were washed for two cycles, resuspended in 100 μL assay buffer, and MFI measured on a FLEXMAP-3D for a minimum of 50 events/region. Bar graphs or binding curves (non-linear 4PL curve) were generated in GraphPad Prism 10.

Assays to examine the interaction of 6His-VHH Chromatibody (137) with biotinylated PTM-defined nucleosomes (the Targets) in the multiplex acidic patch assessment Luminex panel were performed in VHH buffer (20 mM Tris pH 7.5, 100 mM NaCl, 0.01% BSA, 0.01% NP-40, 1 mM DTT) (63). Serially diluted VHH Query (two-fold from 2 μM to 15.6 nM) was probed against the panel (1,000 beads/region per well) following steps above. For detection, 100 μL PE-labeled anti-VHH (1:400; *Jackson ImmunoResearch Laboratories* 128-115-232) was added to each well and incubated for 30 mins with shaking. Beads were washed, resuspended in 100 μL in VHH buffer, and MFI measured. Bar graphs or binding curves (non-linear 4PL curve) were generated in GraphPad Prism 10.

### Native Top-Down Mass Spectrometry (nTDMS)

GST-RAG2 PHD (5 μM) was incubated with nucleosomes (unmodified and/or ([H3K4me3]); 1 μM) in binding buffer (20 mM HEPES pH 7.5, 250 mM NaCl, 0.01% (w/v) BSA, 0.01% (v/v) NP-40, 1 mM DTT) for one hr at 4°C. Reactions were desalted and buffer exchanged to 150 mM Ammonium acetate (AmAc) using a 30 kDa MWCO centrifugal filter (*MilliporeSigma*). Buffer exchange was performed at 10,000 x g for 10 mins at 4 °C and repeated up to 10 times. Protein complex mixtures were adjusted to 1 µM prior to Native MS analysis. Samples were loaded to borosilicate glass emitter tips (*Thermo Fisher Scientific*), Native MS analysis performed on an Ultra High Mass Range (UHMR) Orbitrap QExactive Mass Spectrometer (*Thermo Fisher Scientific*) and data acquired using Xcalibur QualBrowser (*Thermo Fisher Scientific*). First, RAG2-nucleosome complexes were analyzed in MS^1^. Complexes of charge states z = 34+ and 35+ between 8000-8300 m/z range were then activated by collision with nitrogen gas, resulting in ejection and detection of obtained histones by MS^2^. Additional procedures are described elsewhere (93,94,164). MS^1^ and MS^2^ spectra were obtained from averaged scans within MS.raw files; then deconvoluted using UniDec.

### CUT&RUN assays

CUT&RUN (165,166) was performed with 500,000 native K562 cells immobilized on Concanavalin A magnetic beads with a SNAP-CUTANA K-MetStat Panel spike-in (*EpiCypher* 19-1002) to monitor antibody / reader binding. Cells were permeabilized with 0.01% Digitonin and incubated overnight at 4 °C with 0.5 μg antibody (**Suppl. Table 1**) or 70 nM GST-tagged reader (**Suppl. Table 1**). For Reader CUT&RUN reactions, a secondary antibody (0.5 μg anti-GST; *Fortis* A190-122A) was then added. After the addition and activation of pAG-MNase (*EpiCypher* 15-1016), CUT&RUN-enriched DNA was purified using a 2:1 ratio of SPRIselect beads (*EpiCypher* 21-1403). 5 ng DNA was used to prepare sequencing libraries with the CUTANA CUT&RUN Library Prep kit (*EpiCypher* 14-1001). Libraries were analyzed by Agilent TapeStation, pooled to equivalence, and sequenced (*Illumina* NexSeq 2000), targeting ~10 million paired-end reads per reaction.

### Genomic Data Analysis

Paired-end FASTQ files were aligned to the T2T-CHM13v2.0 reference genome using Bowtie2 (167,168). DAC Exclusion List Regions and multi-aligned or duplicate reads were removed prior to subsequent analyses using SAMtools v1.6 and Picard v2.26.2 (169). RPKM-normalized smoothed bigWig files were generated using deepTOOLS v3.5.1 (170) and visualized using Integrative Genome Viewer (IGV). Peaks were called using SICER2 v1.0.3 (171). Peak overlap (≥ 50%) was determined using Intervene v0.6.5 (172) and visualized using matplotlib-venn v0.11.7. deepTools was used to generate TSS and gene body heatmaps and perform Pearson correlations. Antibody and reader domain preference in each reaction was determined by comparing the number of sequencing reads for each DNA barcode in the K-MetStat Panel. anti-PTM and reader domain barcode counts were normalized to reads from the anticipated Target; anti-IgG and anti-GST controls were normalized to total barcode read counts. All sequencing data is available at NCBI Gene Expression Omnibus (accession number GSE249239 / access token: utopgsycpfsbjod). CUT&RUN analyses were performed independently a minimum of three times with consistent results.

K562 RNA-seq data (two replicates of polyA-mRNA) was downloaded from ENCODE (SRR4235541, SRR4235542) (173) and aligned to T2T-CHM13v2.0 using the GTEx RNAseq pipeline (<u>github.com/broadinstitute/gtex-pipeline</u>). Gene expression values were FPKM normalized and replicates assessed for reproducibility (R^2^ = 0.87). Non-protein coding and transcriptionally inactive genes (FPKM = 0) were removed from further analysis and the remaining 15,568 genes binned to deciles by average FPKM. deepTools (170) was used to visualize CUT&RUN signal by RNA-seq decile.

## AUTHOR CONTRIBUTIONS

M.R.M., I.K.P., M.-C.K., and J.M.B. conceptualized and designed the study. M.R.M. and I.K.P. led Captify-Alpha and Captify-Luminex development and experiments using PTM-defined peptides and nucleosomes (conceptualized and/or synthesized by A.V., J.R.B., B.A.B., P.J.B., R.J.E., E.G., Z.B.G., S.A.H., K.K., E.F.P., L.S., H.F.T., R.W., M.A.C., M.J.M., Z.-W.S., and J.M.B.). N.W.H., A.V., J.R.B, T.M.F., S.L.G., N.L.H., V.T.H., A.L.J., L.F.K., K.E.M., K.E.N., K.L.R., C.E.S., L.S., H.E.W., M.J.M., M.A.C, Z.-W.S., M.W.C., E.N.W., M.C.-K., and J.M.B. contributed to Captify assay development, validation and data analysis. S.L.G., N.L.H. and K.E.M. developed and tested VHH domains and analyzed Captify data. Reader-CUT&RUN was performed by E.T.M, D.N.M. and B.J.V., with data analysis and visualization by A.R.H.. Reader domains / constructs were provided by P.J.B., H.A.F., E.G., J.C.S., and C.A.M.. A.S.L., L.F.S. and N.L.K. performed and analyzed RAG2 mass spectrometry studies. M.R.M., E.N.W., M.-C.K. and J.M.B. led writing of the manuscript, with all authors contributing data interpretation and text edits.

## Supporting information

Supplemental Table 1 - Resources

Supplemental Table 2 - Captify Data

## ACKNOWLEDGEMENTS & FUNDING

The authors thank Dr. Masoud Vedadi (Ontario Institute for Cancer Research) for reagent support. The Structural Genomics Consortium (SGC) is a registered charity (No. 1097737) that receives funds from Bayer AG, Boehringer Ingelheim, Bristol Myers Squibb, EU/EFPIA/OICR/McGill/KTH/Diamond Innovative Medicines Initiative 2 Joint Undertaking [EUbOPEN grant 875510], Genentech, Genome Canada through the Ontario Genomics Institute [OGI-196], Janssen, Merck KGaA (*aka* EMD in Canada and USA), Pfizer and Takeda. This work was supported by the National Institutes of Health (NIH) through grant P41GM108569 for the National Resource for Translational and Developmental Proteomics at Northwestern University. A.S.L is a trainee fellow under the Chemistry of Life Processes Predoctoral Training Grant (5T32GM105538-10) at Northwestern University. Work in the Musselman laboratory is funded by NIH grant R35GM128705. *EpiCypher* is supported by NIH grants R43GM134834, R44GM117683, R44GM145007, R44HG010595, R44CA214076, R44GM116584, R44GM119893 and R44HG010640.

## ABBREVIATIONS

ATL: AT-hook-like
BD: bromodomain
CAP: chromatin-associated protein
CD: chromodomain
dNuc: designer nucleosome
MLA: methyl-lysine analog
nTDMS: native top-down mass spectrometry
PHD: plant homeodomain
PTM: post-translational modification
salDNA: salmon sperm DNA
S/B: signal-to-background

## REFERENCES

1. Brownell, J.E., Zhou, J., Ranalli, T., Kobayashi, R., Edmondson, D.G., Roth, S.Y. and Allis, C.D. (1996) Tetrahymena histone acetyltransferase A: a homolog to yeast Gcn5p linking histone acetylation to gene activation. Cell, 84, 843–851.

2. Taunton, J., Hassig, C.A. and Schreiber, S.L. (1996) A mammalian histone deacetylase related to the yeast transcriptional regulator Rpd3p. Science, 272, 408–411.

3. Utley, R.T., Ikeda, K., Grant, P.A., Cote, J., Steger, D.J., Eberharter, A., John, S. and Workman, J.L. (1998) Transcriptional activators direct histone acetyltransferase complexes to nucleosomes. Nature, 394, 498–502.

4. Li, B., Gogol, M., Carey, M., Lee, D., Seidel, C. and Workman, J.L. (2007) Combined action of PHD and chromo domains directs the Rpd3S HDAC to transcribed chromatin. Science, 316, 1050–1054.

5. van Attikum, H. and Gasser, S.M. (2005) The histone code at DNA breaks: a guide to repair? Nat Rev Mol Cell Biol, 6, 757–765.

6. Soria, G., Polo, S.E. and Almouzni, G. (2012) Prime, repair, restore: the active role of chromatin in the DNA damage response. Mol Cell, 46, 722–734.

7. Lee, J.H., Hart, S.R. and Skalnik, D.G. (2004) Histone deacetylase activity is required for embryonic stem cell differentiation. Genesis, 38, 32–38.

8. Ballas, N., Grunseich, C., Lu, D.D., Speh, J.C. and Mandel, G. (2005) REST and its corepressors mediate plasticity of neuronal gene chromatin throughout neurogenesis. Cell, 121, 645–657.

9. Meshorer, E. and Misteli, T. (2006) Chromatin in pluripotent embryonic stem cells and differentiation. Nat Rev Mol Cell Biol, 7, 540–546.

10. Mirabella, A.C., Foster, B.M. and Bartke, T. (2016) Chromatin deregulation in disease. Chromosoma, 125, 75–93.

11. Valencia, A.M. and Kadoch, C. (2019) Chromatin regulatory mechanisms and therapeutic opportunities in cancer. Nat Cell Biol, 21, 152–161.

12. Portela, A. and Esteller, M. (2010) Epigenetic modifications and human disease. Nat Biotechnol, 28, 1057–1068.

13. Luger, K., Mader, A.W., Richmond, R.K., Sargent, D.F. and Richmond, T.J. (1997) Crystal structure of the nucleosome core particle at 2.8 A resolution. Nature, 389, 251–260.

14. Taverna, S.D., Li, H., Ruthenburg, A.J., Allis, C.D. and Patel, D.J. (2007) How chromatin-binding modules interpret histone modifications: lessons from professional pocket pickers. Nat Struct Mol Biol, 14, 1025–1040.

15. Millan-Zambrano, G., Burton, A., Bannister, A.J. and Schneider, R. (2022) Histone post-translational modifications - cause and consequence of genome function. Nat Rev Genet, 23, 563–580.

16. Su, Z. and Denu, J.M. (2016) Reading the Combinatorial Histone Language. ACS Chem Biol, 11, 564–574.

17. Strahl, B.D. and Allis, C.D. (2000) The language of covalent histone modifications. Nature, 403, 41–45.

18. Jenuwein, T. and Allis, C.D. (2001) Translating the histone code. Science, 293, 1074–1080.

19. Lee, J.S., Smith, E. and Shilatifard, A. (2010) The language of histone crosstalk. Cell, 142, 682–685.

20. Smith, E. and Shilatifard, A. (2010) The chromatin signaling pathway: diverse mechanisms of recruitment of histone-modifying enzymes and varied biological outcomes. Mol Cell, 40, 689–701.

21. Henikoff, S. and Shilatifard, A. (2011) Histone modification: cause or cog? Trends Genet, 27, 389–396.

22. Musselman, C.A., Lalonde, M.E., Cote, J. and Kutateladze, T.G. (2012) Perceiving the epigenetic landscape through histone readers. Nat Struct Mol Biol, 19, 1218–1227.

23. Allis, C.D. and Jenuwein, T. (2016) The molecular hallmarks of epigenetic control. Nat Rev Genet, 17, 487–500.

24. Weinzapfel, E.N., Fedder-Semmes, K.N., Sun, Z.W. and Keogh, M.C. (2024) Beyond the tail: the consequence of context in histone post-translational modification and chromatin research. Biochem J, 481, 219–244.

25. Biswas, S. and Rao, C.M. (2018) Epigenetic tools (The Writers, The Readers and The Erasers) and their implications in cancer therapy. Eur J Pharmacol, 837, 8–24.

26. Muller, M.M. and Muir, T.W. (2015) Histones: at the crossroads of peptide and protein chemistry. Chem Rev, 115, 2296–2349.

27. Marunde, M.R., Fuchs, H.A., Burg, J.M., Popova, I.K., Vaidya, A., Hall, N.W., Weinzapfel, E.N., Meiners, M.J., Watson, R., Gillespie, Z.B. et al. (2024) Nucleosome conformation dictates the histone code. Elife, 13, e78866.

28. Marunde, M.R., Popova, I.K., Weinzapfel, E.N. and Keogh, M.C. (2022) The dCypher Approach to Interrogate Chromatin Reader Activity Against Posttranslational Modification-Defined Histone Peptides and Nucleosomes. Methods Mol Biol, 2458, 231–255.

29. Rothbart, S.B., Krajewski, K., Strahl, B.D. and Fuchs, S.M. (2012) Peptide microarrays to interrogate the “histone code”. Methods Enzymol, 512, 107–135.

30. Bua, D.J., Kuo, A.J., Cheung, P., Liu, C.L., Migliori, V., Espejo, A., Casadio, F., Bassi, C., Amati, B., Bedford, M.T. et al. (2009) Epigenome microarray platform for proteome-wide dissection of chromatin-signaling networks. PLoS One, 4, e6789.

31. Filippakopoulos, P., Picaud, S., Mangos, M., Keates, T., Lambert, J.P., Barsyte-Lovejoy, D., Felletar, I., Volkmer, R., Muller, S., Pawson, T. et al. (2012) Histone recognition and large-scale structural analysis of the human bromodomain family. Cell, 149, 214–231.

32. Shah, R.N., Grzybowski, A.T., Cornett, E.M., Johnstone, A.L., Dickson, B.M., Boone, B.A., Cheek, M.A., Cowles, M.W., Maryanski, D., Meiners, M.J. et al. (2018) Examining the Roles of H3K4 Methylation States with Systematically Characterized Antibodies. Mol Cell, 72, 162–177 e167.

33. Szymczak, L.C., Kuo, H.Y. and Mrksich, M. (2018) Peptide Arrays: Development and Application. Anal Chem, 90, 266–282.

34. Petell, C.J., Pham, A.T., Skela, J. and Strahl, B.D. (2019) Improved methods for the detection of histone interactions with peptide microarrays. Sci Rep, 9, 6265.

35. Fang, J., Feng, Q., Ketel, C.S., Wang, H., Cao, R., Xia, L., Erdjument-Bromage, H., Tempst, P., Simon, J.A. and Zhang, Y. (2002) Purification and functional characterization of SET8, a nucleosomal histone H4-lysine 20-specific methyltransferase. Curr Biol, 12, 1086–1099.

36. Li, Y., Trojer, P., Xu, C.F., Cheung, P., Kuo, A., Drury, W.J., 3rd, Qiao, Q., Neubert, T.A., Xu, R.M., Gozani, O. et al. (2009) The target of the NSD family of histone lysine methyltransferases depends on the nature of the substrate. J Biol Chem, 284, 34283–34295.

37. Sankaran, S.M., Wilkinson, A.W., Elias, J.E. and Gozani, O. (2016) A PWWP Domain of Histone-Lysine N-Methyltransferase NSD2 Binds to Dimethylated Lys-36 of Histone H3 and Regulates NSD2 Function at Chromatin. J Biol Chem, 291, 8465–8474.

38. Shi, Y., Lan, F., Matson, C., Mulligan, P., Whetstine, J.R., Cole, P.A., Casero, R.A. and Shi, Y. (2004) Histone demethylation mediated by the nuclear amine oxidase homolog LSD1. Cell, 119, 941–953.

39. Chen, Y., Yang, Y., Wang, F., Wan, K., Yamane, K., Zhang, Y. and Lei, M. (2006) Crystal structure of human histone lysine-specific demethylase 1 (LSD1). Proc Natl Acad Sci U S A, 103, 13956–13961.

40. Yang, M., Gocke, C.B., Luo, X., Borek, D., Tomchick, D.R., Machius, M., Otwinowski, Z. and Yu, H. (2006) Structural basis for CoREST-dependent demethylation of nucleosomes by the human LSD1 histone demethylase. Mol Cell, 23, 377–387.

41. Feng, Q., Wang, H., Ng, H.H., Erdjument-Bromage, H., Tempst, P., Struhl, K. and Zhang, Y. (2002) Methylation of H3-lysine 79 is mediated by a new family of HMTases without a SET domain. Curr Biol, 12, 1052–1058.

42. Min, J., Feng, Q., Li, Z., Zhang, Y. and Xu, R.M. (2003) Structure of the catalytic domain of human DOT1L, a non-SET domain nucleosomal histone methyltransferase. Cell, 112, 711–723.

43. Kuzmichev, A., Nishioka, K., Erdjument-Bromage, H., Tempst, P. and Reinberg, D. (2002) Histone methyltransferase activity associated with a human multiprotein complex containing the Enhancer of Zeste protein. Genes Dev, 16, 2893–2905.

44. Cao, R., Wang, L., Wang, H., Xia, L., Erdjument-Bromage, H., Tempst, P., Jones, R.S. and Zhang, Y. (2002) Role of histone H3 lysine 27 methylation in Polycomb-group silencing. Science, 298, 1039–1043.

45. Czermin, B., Melfi, R., McCabe, D., Seitz, V., Imhof, A. and Pirrotta, V. (2002) Drosophila enhancer of Zeste/ESC complexes have a histone H3 methyltransferase activity that marks chromosomal Polycomb sites. Cell, 111, 185–196.

46. Thomas, J.F., Valencia-Sanchez, M.I., Tamburri, S., Gloor, S.L., Rustichelli, S., Godinez-Lopez, V., De Ioannes, P., Lee, R., Abini-Agbomson, S., Gretarsson, K., et al. (2023) Structural basis of histone H2A lysine 119 deubiquitination by Polycomb Repressive Deubiquitinase BAP1/ASXL1. Sci Adv., 9, eadg9832.

47. Morrison, E.A., Bowerman, S., Sylvers, K.L., Wereszczynski, J. and Musselman, C.A. (2018) The conformation of the histone H3 tail inhibits association of the BPTF PHD finger with the nucleosome. Elife, 7, e31481.

48. Weaver, T.M., Morrison, E.A. and Musselman, C.A. (2018) Reading More than Histones: The Prevalence of Nucleic Acid Binding among Reader Domains. Molecules, 23, 2614.

49. Kasinath, V., Beck, C., Sauer, P., Poepsel, S., Kosmatka, J., Faini, M., Toso, D., Aebersold, R. and Nogales, E. (2021) JARID2 and AEBP2 regulate PRC2 in the presence of H2AK119ub1 and other histone modifications. Science, 371, eabc3393.

50. 50. Valencia-Sanchez, M.I., De Ioannes, P., Wang, M., Vasilyev, N., Chen, R., Nudler, E., Armache, J.P. and Armache, K.J. (2019) Structural Basis of Dot1L Stimulation by Histone H2B Lysine 120 Ubiquitination. Mol Cell, 74, 1010–1019.e1016.

51. Jang, S., Kang, C., Yang, H.S., Jung, T., Hebert, H., Chung, K.Y., Kim, S.J., Hohng, S. and Song, J.J. (2019) Structural basis of recognition and destabilization of the histone H2B ubiquitinated nucleosome by the DOT1L histone H3 Lys79 methyltransferase. Genes Dev, 33, 620–625.

52. Worden, E.J., Hoffmann, N.A., Hicks, C.W. and Wolberger, C. (2019) Mechanism of Cross-talk between H2B Ubiquitination and H3 Methylation by Dot1L. Cell, 176, 1490–1501 e1412.

53. Anderson, C.J., Baird, M.R., Hsu, A., Barbour, E.H., Koyama, Y., Borgnia, M.J. and McGinty, R.K. (2019) Structural Basis for Recognition of Ubiquitylated Nucleosome by Dot1L Methyltransferase. Cell Rep, 26, 1681–1690 e1685.

54. Vaughan, R.M., Dickson, B.M., Whelihan, M.F., Johnstone, A.L., Cornett, E.M., Cheek, M.A., Ausherman, C.A., Cowles, M.W., Sun, Z.W. and Rothbart, S.B. (2018) Chromatin structure and its chemical modifications regulate the ubiquitin ligase substrate selectivity of UHRF1. Proc Natl Acad Sci U S A, 115, 8775–8780.

55. Jain, K., Marunde, M.R., Burg, J.M., Gloor, S.L., Joseph, F.M., Poncha, K.F., Gillespie, Z.B., Rodriguez, K.L., Popova, I.K., Hall, N.W. et al. (2023) An acetylation-mediated chromatin switch governs H3K4 methylation read-write capability. Elife, 12, e82596.

56. Keogh, M.-C., Almouzni, G., Andrews, A.J., Armache, K.-J., Arrowsmith, C.H., Baek, S.H., Bedford, M.T., Bernstein, E., David, Y., Denu, J.M. et al. A Needed Nomenclature for Nucleosomes. Manuscript under review.

57. Dilworth, D., Hanley, R.P., Ferreira de Freitas, R., Allali-Hassani, A., Zhou, M., Mehta, N., Marunde, M.R., Ackloo, S., Carvalho Machado, R.A., Khalili Yazdi, A., et al. (2022) A chemical probe targeting the PWWP domain alters NSD2 nucleolar localization. Nat Chem Biol, 18, 56–63.

58. Hanley, R.P., Nie, D.Y., Tabor, J.R., Li, F., Sobh, A., Xu, C., Barker, N.K., Dilworth, D., Hajian, T., Gibson, E. et al. (2023) Discovery of a Potent and Selective Targeted NSD2 Degrader for the Reduction of H3K36me2. J Am Chem Soc, 145, 8176–8188.

59. Morgan, M.A.J., Popova, I.K., Vaidya, A., Burg, J.M., Marunde, M.R., Rendleman, E.J., Dumar, Z.J., Watson, R., Meiners, M.J., Howard, S.A. et al. (2021) A trivalent nucleosome interaction by PHIP/BRWD2 is disrupted in neurodevelopmental disorders and cancer. Genes Dev, 35, 1642–1656.

60. Weinberg, D.N., Papillon-Cavanagh, S., Chen, H., Yue, Y., Chen, X., Rajagopalan, K.N., Horth, C., McGuire, J.T., Xu, X., Nikbakht, H. et al. (2019) The histone mark H3K36me2 recruits DNMT3A and shapes the intergenic DNA methylation landscape. Nature, 573, 281–286.

61. Weinberg, D.N., Rosenbaum, P., Chen, X., Barrows, D., Horth, C., Marunde, M.R., Popova, I.K., Gillespie, Z.B., Keogh, M.C., Lu, C. et al. (2021) Two competing mechanisms of DNMT3A recruitment regulate the dynamics of de novo DNA methylation at PRC1-targeted CpG islands. Nat Genet, 53, 794–800.

62. Gretarsson, K.H., Abini-Agbomson, S., Gloor, S.L., Weinberg, D.N., McCuiston, J.L., Kumary, V.U.S., Hickman, A.R., Sahu, V., Lee, R., Xu, X. et al. (2024) Cancer-associated DNA Hypermethylation of Polycomb Targets Requires DNMT3A Dual Recognition of Histone H2AK119 Ubiquitination and the Nucleosome Acidic Patch. Science advances, 10, eadp0975.

63. Hicks, C.W., Rahman, S., Gloor, S.L., Fields, J.K., Husby, N.L., Vaidya, A., Maier, K.E., Morgan, M., Keogh, M.C. and Wolberger, C. (2024) Ubiquitinated histone H2B as gatekeeper of the nucleosome acidic patch. Nucleic Acids Res, 52, 9978–9995.

64. Eglen, R.M., Reisine, T., Roby, P., Rouleau, N., Illy, C., Bosse, R. and Bielefeld, M. (2008) The use of AlphaScreen technology in HTS: current status. Curr Chem Genomics, 1, 2–10.

65. Jung, M., Philpott, M., Muller, S., Schulze, J., Badock, V., Eberspacher, U., Moosmayer, D., Bader, B., Schmees, N., Fernandez-Montalvan, A. et al. (2014) Affinity map of bromodomain protein 4 (BRD4) interactions with the histone H4 tail and the small molecule inhibitor JQ1. J Biol Chem, 289, 9304–9319.

66. Holt, M. and Muir, T. (2015) Application of the protein semisynthesis strategy to the generation of modified chromatin. Annu Rev Biochem, 84, 265–290.

67. 67. (2004) In Markossian, S., Grossman, A., Baskir, H., Arkin, M., Auld, D., Austin, C., Baell, J., Brimacombe, K., Chung, T. D. Y., Coussens, N. P. et al. (eds.), Assay Guidance Manual, Bethesda (MD).

68. Varela, I., Tarpey, P., Raine, K., Huang, D., Ong, C.K., Stephens, P., Davies, H., Jones, D., Lin, M.L., Teague, J. et al. (2011) Exome sequencing identifies frequent mutation of the SWI/SNF complex gene PBRM1 in renal carcinoma. Nature, 469, 539–542.

69. Morrison, E.A., Sanchez, J.C., Ronan, J.L., Farrell, D.P., Varzavand, K., Johnson, J.K., Gu, B.X., Crabtree, G.R. and Musselman, C.A. (2017) DNA binding drives the association of BRG1/hBRM bromodomains with nucleosomes. Nat Commun, 8, 16080.

70. Agalioti, T., Chen, G. and Thanos, D. (2002) Deciphering the transcriptional histone acetylation code for a human gene. Cell, 111, 381–392.

71. Schulze, J.M., Wang, A.Y. and Kobor, M.S. (2009) YEATS domain proteins: a diverse family with many links to chromatin modification and transcription. Biochem Cell Biol, 87, 65–75.

72. Zhao, D., Li, Y., Xiong, X., Chen, Z. and Li, H. (2017) YEATS Domain-A Histone Acylation Reader in Health and Disease. J Mol Biol, 429, 1994–2002.

73. Yeewa, R., Chaiya, P., Jantrapirom, S., Shotelersuk, V. and Lo Piccolo, L. (2022) Multifaceted roles of YEATS domain-containing proteins and novel links to neurological diseases. Cell Mol Life Sci, 79, 183.

74. Allis, C.D., Berger, S.L., Cote, J., Dent, S., Jenuwien, T., Kouzarides, T., Pillus, L., Reinberg, D., Shi, Y., Shiekhattar, R. et al. (2007) New nomenclature for chromatin-modifying enzymes. Cell, 131, 633–636.

75. Tian, X., Zhang, S., Liu, H.M., Zhang, Y.B., Blair, C.A., Mercola, D., Sassone-Corsi, P. and Zi, X. (2013) Histone lysine-specific methyltransferases and demethylases in carcinogenesis: new targets for cancer therapy and prevention. Curr Cancer Drug Targets, 13, 558–579.

76. Hyun, K., Jeon, J., Park, K. and Kim, J. (2017) Writing, erasing and reading histone lysine methylations. Exp Mol Med, 49, e324.

77. Husmann, D. and Gozani, O. (2019) Histone lysine methyltransferases in biology and disease. Nat Struct Mol Biol, 26, 880–889.

78. Bhat, K.P., Umit Kaniskan, H., Jin, J. and Gozani, O. (2021) Epigenetics and beyond: targeting writers of protein lysine methylation to treat disease. Nat Rev Drug Discov, 20, 265–286.

79. Kalakonda, N., Fischle, W., Boccuni, P., Gurvich, N., Hoya-Arias, R., Zhao, X., Miyata, Y., Macgrogan, D., Zhang, J., Sims, J.K. et al. (2008) Histone H4 lysine 20 monomethylation promotes transcriptional repression by L3MBTL1. Oncogene, 27, 4293–4304.

80. Kim, J., Daniel, J., Espejo, A., Lake, A., Krishna, M., Xia, L., Zhang, Y. and Bedford, M.T. (2006) Tudor, MBT and chromo domains gauge the degree of lysine methylation. EMBO Rep, 7, 397–403.

81. Trojer, P., Li, G., Sims, R.J., 3rd, Vaquero, A., Kalakonda, N., Boccuni, P., Lee, D., Erdjument-Bromage, H., Tempst, P., Nimer, S.D. et al. (2007) L3MBTL1, a histone-methylation-dependent chromatin lock. Cell, 129, 915–928.

82. Li, H., Fischle, W., Wang, W., Duncan, E.M., Liang, L., Murakami-Ishibe, S., Allis, C.D. and Patel, D.J. (2007) Structural basis for lower lysine methylation state-specific readout by MBT repeats of L3MBTL1 and an engineered PHD finger. Mol Cell, 28, 677–691.

83. Min, J., Allali-Hassani, A., Nady, N., Qi, C., Ouyang, H., Liu, Y., MacKenzie, F., Vedadi, M. and Arrowsmith, C.H. (2007) L3MBTL1 recognition of mono- and dimethylated histones. Nat Struct Mol Biol, 14, 1229–1230.

84. Sbardella, G. (2020) In Mai, A. (ed.), Chemical Epigenetics. Springer International Publishing, Cham, pp. 339–399.

85. MacGrogan, D., Kalakonda, N., Alvarez, S., Scandura, J.M., Boccuni, P., Johansson, B. and Nimer, S.D. (2004) Structural integrity and expression of the L3MBTL gene in normal and malignant hematopoietic cells. Genes Chromosomes Cancer, 41, 203–213.

86. Gurvich, N., Perna, F., Farina, A., Voza, F., Menendez, S., Hurwitz, J. and Nimer, S.D. (2010) L3MBTL1 polycomb protein, a candidate tumor suppressor in del(20q12) myeloid disorders, is essential for genome stability. Proc Natl Acad Sci U S A, 107, 22552–22557.

87. Oettinger, M.A., Schatz, D.G., Gorka, C. and Baltimore, D. (1990) RAG-1 and RAG-2, adjacent genes that synergistically activate V(D)J recombination. Science, 248, 1517–1523.

88. Schatz, D.G. and Swanson, P.C. (2011) V(D)J recombination: mechanisms of initiation. Annu Rev Genet, 45, 167–202.

89. Qiu, J.X., Kale, S.B., Yarnell Schultz, H. and Roth, D.B. (2001) Separation-of-function mutants reveal critical roles for RAG2 in both the cleavage and joining steps of V(D)J recombination. Mol Cell, 7, 77–87.

90. Matthews, A.G., Kuo, A.J., Ramon-Maiques, S., Han, S., Champagne, K.S., Ivanov, D., Gallardo, M., Carney, D., Cheung, P., Ciccone, D.N. et al. (2007) RAG2 PHD finger couples histone H3 lysine 4 trimethylation with V(D)J recombination. Nature, 450, 1106–1110.

91. Ramon-Maiques, S., Kuo, A.J., Carney, D., Matthews, A.G., Oettinger, M.A., Gozani, O. and Yang, W. (2007) The plant homeodomain finger of RAG2 recognizes histone H3 methylated at both lysine-4 and arginine-2. Proc Natl Acad Sci U S A, 104, 18993–18998.

92. Liu, Y., Subrahmanyam, R., Chakraborty, T., Sen, R. and Desiderio, S. (2007) A plant homeodomain in RAG-2 that binds Hypermethylated lysine 4 of histone H3 is necessary for efficient antigen-receptor-gene rearrangement. Immunity, 27, 561–571.

93. Schachner, L.F., Jooss, K., Morgan, M.A., Piunti, A., Meiners, M.J., Kafader, J.O., Lee, A.S., Iwanaszko, M., Cheek, M.A., Burg, J.M. et al. (2021) Decoding the protein composition of whole nucleosomes with Nuc-MS. Nat Methods, 18, 303–308.

94. Jooss, K., Schachner, L.F., Watson, R., Gillespie, Z.B., Howard, S.A., Cheek, M.A., Meiners, M.J., Sobh, A., Licht, J.D., Keogh, M.C. et al. (2021) Separation and Characterization of Endogenous Nucleosomes by Native Capillary Zone Electrophoresis-Top-Down Mass Spectrometry. Anal Chem, 93, 5151–5160.

95. Nakayama, J., Rice, J.C., Strahl, B.D., Allis, C.D. and Grewal, S.I. (2001) Role of histone H3 lysine 9 methylation in epigenetic control of heterochromatin assembly. Science, 292, 110–113.

96. Bannister, A.J., Zegerman, P., Partridge, J.F., Miska, E.A., Thomas, J.O., Allshire, R.C. and Kouzarides, T. (2001) Selective recognition of methylated lysine 9 on histone H3 by the HP1 chromo domain. Nature, 410, 120–124.

97. Lachner, M., O’Carroll, D., Rea, S., Mechtler, K. and Jenuwein, T. (2001) Methylation of histone H3 lysine 9 creates a binding site for HP1 proteins. Nature, 410, 116–120.

98. Bernstein, E., Duncan, E.M., Masui, O., Gil, J., Heard, E. and Allis, C.D. (2006) Mouse polycomb proteins bind differentially to methylated histone H3 and RNA and are enriched in facultative heterochromatin. Mol Cell Biol, 26, 2560–2569.

99. Tardat, M., Albert, M., Kunzmann, R., Liu, Z., Kaustov, L., Thierry, R., Duan, S., Brykczynska, U., Arrowsmith, C.H. and Peters, A.H. (2015) Cbx2 targets PRC1 to constitutive heterochromatin in mouse zygotes in a parent-of-origin-dependent manner. Mol Cell, 58, 157–171.

100. Kaustov, L., Ouyang, H., Amaya, M., Lemak, A., Nady, N., Duan, S., Wasney, G.A., Li, Z., Vedadi, M., Schapira, M. et al. (2011) Recognition and specificity determinants of the human cbx chromodomains. J Biol Chem, 286, 521–529.

101. Zhen, C.Y., Tatavosian, R., Huynh, T.N., Duc, H.N., Das, R., Kokotovic, M., Grimm, J.B., Lavis, L.D., Lee, J., Mejia, F.J. et al. (2016) Live-cell single-molecule tracking reveals co-recognition of H3K27me3 and DNA targets polycomb Cbx7-PRC1 to chromatin. Elife, 5, e17667.

102. Senthilkumar, R. and Mishra, R.K. (2009) Novel motifs distinguish multiple homologues of Polycomb in vertebrates: expansion and diversification of the epigenetic toolkit. BMC Genomics, 10, 549.

103. Dhayalan, A., Rajavelu, A., Rathert, P., Tamas, R., Jurkowska, R.Z., Ragozin, S. and Jeltsch, A. (2010) The Dnmt3a PWWP domain reads histone 3 lysine 36 trimethylation and guides DNA methylation. J Biol Chem, 285, 26114–26120.

104. Vermeulen, M., Eberl, H.C., Matarese, F., Marks, H., Denissov, S., Butter, F., Lee, K.K., Olsen, J.V., Hyman, A.A., Stunnenberg, H.G. et al. (2010) Quantitative interaction proteomics and genome-wide profiling of epigenetic histone marks and their readers. Cell, 142, 967–980.

105. Wu, H., Zeng, H., Lam, R., Tempel, W., Amaya, M.F., Xu, C., Dombrovski, L., Qiu, W., Wang, Y. and Min, J. (2011) Structural and histone binding ability characterizations of human PWWP domains. PLoS One, 6, e18919.

106. Iwase, S., Xiang, B., Ghosh, S., Ren, T., Lewis, P.W., Cochrane, J.C., Allis, C.D., Picketts, D.J., Patel, D.J., Li, H. et al. (2011) ATRX ADD domain links an atypical histone methylation recognition mechanism to human mental-retardation syndrome. Nat Struct Mol Biol, 18, 769–776.

107. Eustermann, S., Yang, J.C., Law, M.J., Amos, R., Chapman, L.M., Jelinska, C., Garrick, D., Clynes, D., Gibbons, R.J., Rhodes, D. et al. (2011) Combinatorial readout of histone H3 modifications specifies localization of ATRX to heterochromatin. Nat Struct Mol Biol, 18, 777–782.

108. Simon, M.D., Chu, F., Racki, L.R., de la Cruz, C.C., Burlingame, A.L., Panning, B., Narlikar, G.J. and Shokat, K.M. (2007) The site-specific installation of methyl-lysine analogs into recombinant histones. Cell, 128, 1003–1012.

109. Simon, M.D. (2010) Installation of site-specific methylation into histones using methyl lysine analogs. Curr Protoc Mol Biol, Chapter 21, Unit 21 18 21-10.

110. Simon, M.D. and Shokat, K.M. (2012) A method to site-specifically incorporate methyl-lysine analogues into recombinant proteins. Methods Enzymol, 512, 57–69.

111. Chen, Z., Grzybowski, A.T. and Ruthenburg, A.J. (2014) Traceless semisynthesis of a set of histone 3 species bearing specific lysine methylation marks. Chembiochem, 15, 2071–2075.

112. Chen, Z., Notti, R.Q., Ueberheide, B. and Ruthenburg, A.J. (2018) Quantitative and Structural Assessment of Histone Methyllysine Analogue Engagement by Cognate Binding Proteins Reveals Affinity Decrements Relative to Those of Native Counterparts. Biochemistry, 57, 300–304.

113. Munari, F., Soeroes, S., Zenn, H.M., Schomburg, A., Kost, N., Schroder, S., Klingberg, R., Rezaei-Ghaleh, N., Stutzer, A., Gelato, K.A. et al. (2012) Methylation of lysine 9 in histone H3 directs alternative modes of highly dynamic interaction of heterochromatin protein hHP1beta with the nucleosome. J Biol Chem, 287, 33756–33765.

114. Seeliger, D., Soeroes, S., Klingberg, R., Schwarzer, D., Grubmuller, H. and Fischle, W. (2012) Quantitative assessment of protein interaction with methyl-lysine analogues by hybrid computational and experimental approaches. ACS Chem Biol, 7, 150–154.

115. Ruthenburg, A.J., Li, H., Patel, D.J. and Allis, C.D. (2007) Multivalent engagement of chromatin modifications by linked binding modules. Nat Rev Mol Cell Biol, 8, 983–994.

116. Fischle, W., Tseng, B.S., Dormann, H.L., Ueberheide, B.M., Garcia, B.A., Shabanowitz, J., Hunt, D.F., Funabiki, H. and Allis, C.D. (2005) Regulation of HP1-chromatin binding by histone H3 methylation and phosphorylation. Nature, 438, 1116–1122.

117. Rea, S., Eisenhaber, F., O’Carroll, D., Strahl, B.D., Sun, Z.W., Schmid, M., Opravil, S., Mechtler, K., Ponting, C.P., Allis, C.D. et al. (2000) Regulation of chromatin structure by site-specific histone H3 methyltransferases. Nature, 406, 593–599.

118. Kunowska, N., Rotival, M., Yu, L., Choudhary, J. and Dillon, N. (2015) Identification of protein complexes that bind to histone H3 combinatorial modifications using super-SILAC and weighted correlation network analysis. Nucleic Acids Res, 43, 1418–1432.

119. Noh, K.M., Maze, I., Zhao, D., Xiang, B., Wenderski, W., Lewis, P.W., Shen, L., Li, H. and Allis, C.D. (2015) ATRX tolerates activity-dependent histone H3 methyl/phos switching to maintain repetitive element silencing in neurons. Proc Natl Acad Sci U S A, 112, 6820–6827.

120. Rona, G.B., Eleutherio, E.C.A. and Pinheiro, A.S. (2016) PWWP domains and their modes of sensing DNA and histone methylated lysines. Biophys Rev, 8, 63–74.

121. Fang, R., Chen, F., Dong, Z., Hu, D., Barbera, A.J., Clark, E.A., Fang, J., Yang, Y., Mei, P., Rutenberg, M. et al. (2013) LSD2/KDM1B and its cofactor NPAC/GLYR1 endow a structural and molecular model for regulation of H3K4 demethylation. Mol Cell, 49, 558–570.

122. Marabelli, C., Marrocco, B., Pilotto, S., Chittori, S., Picaud, S., Marchese, S., Ciossani, G., Forneris, F., Filippakopoulos, P., Schoehn, G. et al. (2019) A Tail-Based Mechanism Drives Nucleosome Demethylation by the LSD2/NPAC Multimeric Complex. Cell Rep, 27, 387–399 e387.

123. Fei, J., Ishii, H., Hoeksema, M.A., Meitinger, F., Kassavetis, G.A., Glass, C.K., Ren, B. and Kadonaga, J.T. (2018) NDF, a nucleosome-destabilizing factor that facilitates transcription through nucleosomes. Genes Dev, 32, 682–694.

124. Yu, S., Li, J., Ji, G., Ng, Z.L., Siew, J., Lo, W.N., Ye, Y., Chew, Y.Y., Long, Y.C., Zhang, W. et al. (2022) Npac Is A Co-factor of Histone H3K36me3 and Regulates Transcriptional Elongation in Mouse Embryonic Stem Cells. Genomics Proteomics Bioinformatics, 20, 110–128.

125. 125. Schuettengruber, B., Bourbon, H.M., Di Croce, L. and Cavalli, G. (2017) Genome Regulation by Polycomb and Trithorax: 70 Years and Counting. Cell, 171, 34–57.

126. Vincenz, C. and Kerppola, T.K. (2008) Different polycomb group CBX family proteins associate with distinct regions of chromatin using nonhomologous protein sequences. Proc Natl Acad Sci U S A, 105, 16572–16577.

127. 127. Morey, L., Pascual, G., Cozzuto, L., Roma, G., Wutz, A., Benitah, S.A. and Di Croce, L. (2012) Nonoverlapping functions of the Polycomb group Cbx family of proteins in embryonic stem cells. Cell Stem Cell, 10, 47–62.

128. Morey, L., Aloia, L., Cozzuto, L., Benitah, S.A. and Di Croce, L. (2013) RYBP and Cbx7 define specific biological functions of polycomb complexes in mouse embryonic stem cells. Cell Rep, 3, 60–69.

129. Skrajna, A., Goldfarb, D., Kedziora, K.M., Cousins, E.M., Grant, G.D., Spangler, C.J., Barbour, E.H., Yan, X., Hathaway, N.A., Brown, N.G. et al. (2020) Comprehensive nucleosome interactome screen establishes fundamental principles of nucleosome binding. Nucleic Acids Res, 48, 9415–9432.

130. McGinty, R.K. and Tan, S. (2021) Principles of nucleosome recognition by chromatin factors and enzymes. Curr Opin Struct Biol, 71, 16–26.

131. Reyes, A.A., Marcum, R.D. and He, Y. (2021) Structure and Function of Chromatin Remodelers. J Mol Biol, 433, 166929.

132. Shogren-Knaak, M., Ishii, H., Sun, J.M., Pazin, M.J., Davie, J.R. and Peterson, C.L. (2006) Histone H4-K16 acetylation controls chromatin structure and protein interactions. Science, 311, 844–847.

133. Henry, K.W., Wyce, A., Lo, W.S., Duggan, L.J., Emre, N.C., Kao, C.F., Pillus, L., Shilatifard, A., Osley, M.A. and Berger, S.L. (2003) Transcriptional activation via sequential histone H2B ubiquitylation and deubiquitylation, mediated by SAGA-associated Ubp8. Genes Dev, 17, 2648–2663.

134. Shema-Yaacoby, E., Nikolov, M., Haj-Yahya, M., Siman, P., Allemand, E., Yamaguchi, Y., Muchardt, C., Urlaub, H., Brik, A., Oren, M. et al. (2013) Systematic identification of proteins binding to chromatin-embedded ubiquitylated H2B reveals recruitment of SWI/SNF to regulate transcription. Cell Rep, 4, 601–608.

135. Dann, G.P., Liszczak, G.P., Bagert, J.D., Muller, M.M., Nguyen, U.T.T., Wojcik, F., Brown, Z.Z., Bos, J., Panchenko, T., Pihl, R. et al. (2017) ISWI chromatin remodellers sense nucleosome modifications to determine substrate preference. Nature, 548, 607–611.

136. Nune, M., Morgan, M.T., Connell, Z., McCullough, L., Jbara, M., Sun, H., Brik, A., Formosa, T. and Wolberger, C. (2019) FACT and Ubp10 collaborate to modulate H2B deubiquitination and nucleosome dynamics. Elife, 8, e40988.

137. Jullien, D., Vignard, J., Fedor, Y., Bery, N., Olichon, A., Crozatier, M., Erard, M., Cassard, H., Ducommun, B., Salles, B. et al. (2016) Chromatibody, a novel non-invasive molecular tool to explore and manipulate chromatin in living cells. J Cell Sci, 129, 2673–2683.

138. Vermeulen, M., Mulder, K.W., Denissov, S., Pijnappel, W.W., van Schaik, F.M., Varier, R.A., Baltissen, M.P., Stunnenberg, H.G., Mann, M. and Timmers, H.T. (2007) Selective anchoring of TFIID to nucleosomes by trimethylation of histone H3 lysine 4. Cell, 131, 58–69.

139. Kungulovski, G., Mauser, R., Reinhardt, R. and Jeltsch, A. (2016) Application of recombinant TAF3 PHD domain instead of anti-H3K4me3 antibody. Epigenetics Chromatin, 9, 11.

140. Chiang, K.P., Jensen, M.S., McGinty, R.K. and Muir, T.W. (2009) A semisynthetic strategy to generate phosphorylated and acetylated histone H2B. Chembiochem, 10, 2182–2187.

141. Hananya, N., Koren, S. and Muir, T.W. (2024) Interrogating epigenetic mechanisms with chemically customized chromatin. Nat Rev Genet, 25, 255–271.

142. Fierz, B. and Muir, T.W. (2012) Chromatin as an expansive canvas for chemical biology. Nat Chem Biol, 8, 417–427.

143. Lukauskas, S., Tvardovskiy, A., Nguyen, N.V., Stadler, M., Faull, P., Ravnsborg, T., Ozdemir Aygenli, B., Dornauer, S., Flynn, H., Lindeboom, R.G.H. et al. (2024) Decoding chromatin states by proteomic profiling of nucleosome readers. Nature, 627, 671–679.

144. Whedon, S.D., Lee, K., Wang, Z.A., Zahn, E., Lu, C., Yapa Abeywardana, M., Fairall, L., Nam, E., DuBois-Coyne, S., De Ioannes, P., et al. (2024) Circular Engineered Sortase for Interrogating Histone H3 in Chromatin. J Am Chem Soc, 146, 33914–33927.

145. Ruthenburg, A.J., Li, H., Milne, T.A., Dewell, S., McGinty, R.K., Yuen, M., Ueberheide, B., Dou, Y., Muir, T.W., Patel, D.J. et al. (2011) Recognition of a mononucleosomal histone modification pattern by BPTF via multivalent interactions. Cell, 145, 692–706.

146. Moriniere, J., Rousseaux, S., Steuerwald, U., Soler-Lopez, M., Curtet, S., Vitte, A.L., Govin, J., Gaucher, J., Sadoul, K., Hart, D.J. et al. (2009) Cooperative binding of two acetylation marks on a histone tail by a single bromodomain. Nature, 461, 664–668.

147. Jacobson, R.H., Ladurner, A.G., King, D.S. and Tjian, R. (2000) Structure and function of a human TAFII250 double bromodomain module. Science, 288, 1422–1425.

148. Owen, D.J., Ornaghi, P., Yang, J.C., Lowe, N., Evans, P.R., Ballario, P., Neuhaus, D., Filetici, P. and Travers, A.A. (2000) The structural basis for the recognition of acetylated histone H4 by the bromodomain of histone acetyltransferase gcn5p. EMBO J, 19, 6141–6149.

149. Chahal, S.S., Matthews, H.R. and Bradbury, E.M. (1980) Acetylation of histone H4 and its role in chromatin structure and function. Nature, 287, 76–79.

150. Cary, P.D., Crane-Robinson, C., Bradbury, E.M. and Dixon, G.H. (1982) Effect of acetylation on the binding of N-terminal peptides of histone H4 to DNA. Eur J Biochem, 127, 137–143.

151. Ebralidse, K.K., Grachev, S.A. and Mirzabekov, A.D. (1988) A highly basic histone H4 domain bound to the sharply bent region of nucleosomal DNA. Nature, 331, 365–367.

152. Hong, L., Schroth, G.P., Matthews, H.R., Yau, P. and Bradbury, E.M. (1993) Studies of the DNA binding properties of histone H4 amino terminus. Thermal denaturation studies reveal that acetylation markedly reduces the binding constant of the H4 “tail” to DNA. J Biol Chem, 268, 305–314.

153. Vettese-Dadey, M., Grant, P.A., Hebbes, T.R., Crane-Robinson, C., Allis, C.D. and Workman, J.L. (1996) Acetylation of histone H4 plays a primary role in enhancing transcription factor binding to nucleosomal DNA in vitro. EMBO J, 15, 2508–2518.

154. Ghoneim, M., Fuchs, H.A. and Musselman, C.A. (2021) Histone Tail Conformations: A Fuzzy Affair with DNA. Trends Biochem Sci, 46, 564–578.

155. Kouzarides, T. (2000) Acetylation: a regulatory modification to rival phosphorylation? EMBO J, 19, 1176–1179.

156. Lee, A.S., Marunde, M.R., Su, P., Schachner, L.F., Khan, L.F., Graham, B., Taylor, H.F., Onuoha, U.C., Jooβ, K., Fuchs, H.A., et al. Direct Readout of Multivalent Chromatin Reader-Nucleosome Interactions by Nucleosome Mass Spectrometry. *Manuscript under review*.

157. Bell, M.R., Engleka, M.J., Malik, A. and Strickler, J.E. (2013) To fuse or not to fuse: what is your purpose? Protein Sci, 22, 1466–1477.

158. Albanese, K.I., Krone, M.W., Petell, C.J., Parker, M.M., Strahl, B.D., Brustad, E.M. and Waters, M.L. (2020) Engineered Reader Proteins for Enhanced Detection of Methylated Lysine on Histones. ACS Chem Biol, 15, 103–111.

159. Dias, J.K. and D’Arcy, S. (2025) Beyond the mono-nucleosome. Biochem Soc Trans, 53, BCJ20240452.

160. Liu, Z., Xi, S., McGregor, L.A., Yamatsugu, K., Kawashima, S.A., Sczepanski, J.T. and Kanai, M. (2025) A Method for Constructing Nucleosome Arrays with Spatially Defined Histone PTMs and DNA Damage. Angew Chem Int Ed Engl, e202500162.

161. Lopez, V.G., Valencia-Sanchez, M.I., Abini-Agbomson, S., Thomas, J.F., Lee, R., De Ioannes, P., Sosa, B.A., Armache, J.P. and Armache, K.J. (2024) Read-write mechanisms of H2A ubiquitination by Polycomb repressive complex 1. Nature, 636, 755–761.

162. Mauser, R., Kungulovski, G., Keup, C., Reinhardt, R. and Jeltsch, A. (2017) Application of dual reading domains as novel reagents in chromatin biology reveals a new H3K9me3 and H3K36me2/3 bivalent chromatin state. Epigenetics Chromatin, 10, 45.

163. Barral, A., Pozo, G., Ducrot, L., Papadopoulos, G.L., Sauzet, S., Oldfield, A.J., Cavalli, G. and Dejardin, J. (2022) SETDB1/NSD-dependent H3K9me3/H3K36me3 dual heterochromatin maintains gene expression profiles by bookmarking poised enhancers. Mol Cell, 82, 816–832 e812.

164. Marty, M.T., Baldwin, A.J., Marklund, E.G., Hochberg, G.K., Benesch, J.L. and Robinson, C.V. (2015) Bayesian deconvolution of mass and ion mobility spectra: from binary interactions to polydisperse ensembles. Anal Chem, 87, 4370–4376.

165. Skene, P.J. and Henikoff, S. (2017) An efficient targeted nuclease strategy for high-resolution mapping of DNA binding sites. Elife, 6, e21856.

166. Firestone, T.M., Venters, B.J., Novitzky, K., Albertorio-Sáez, L.M., Barnes, C.A., Fedder-Semmes, K.N., Hall, N.W., Hickman, A.R., Kaderli, M., Windham, C.L., et al. (2024) High-efficiency genomic mapping of chromatin-associated targets with CUT&RUN. bioRxiv, 2024.2012.2003.626419.

167. Langmead, B. and Salzberg, S.L. (2012) Fast gapped-read alignment with Bowtie 2. Nat Methods, 9, 357–359.

168. Nurk, S., Koren, S., Rhie, A., Rautiainen, M., Bzikadze, A.V., Mikheenko, A., Vollger, M.R., Altemose, N., Uralsky, L., Gershman, A. et al. (2022) The complete sequence of a human genome. Science, 376, 44–53.

169. Danecek, P., Bonfield, J.K., Liddle, J., Marshall, J., Ohan, V., Pollard, M.O., Whitwham, A., Keane, T., McCarthy, S.A., Davies, R.M. et al. (2021) Twelve years of SAMtools and BCFtools. Gigascience, 10, giab008.

170. Ramirez, F., Ryan, D.P., Gruning, B., Bhardwaj, V., Kilpert, F., Richter, A.S., Heyne, S., Dundar, F. and Manke, T. (2016) deepTools2: a next generation web server for deep-sequencing data analysis. Nucleic Acids Res, 44, W160–165.

171. Zang, C., Schones, D.E., Zeng, C., Cui, K., Zhao, K. and Peng, W. (2009) A clustering approach for identification of enriched domains from histone modification ChIP-Seq data. Bioinformatics, 25, 1952–1958.

172. Khan, A. and Mathelier, A. (2017) Intervene: a tool for intersection and visualization of multiple gene or genomic region sets. BMC Bioinformatics, 18, 287.

173. Consortium, E.P. (2012) An integrated encyclopedia of DNA elements in the human genome. Nature, 489, 57–74.

