## Supplemental Figure Legends for "Nucleosome context regulates chromatin reader preference"

### SUPPLEMENTARY TABLES

**Suppl. Table 1A-E:** Resources

**Tab A:** Expression Plasmids and Recombinant Proteins

**Tab B:** Captify™ peptides

**Tab C:** Captify dNucs

**Tab D:** versaNucs & related peptides

**Tab E:** Luminex Panels

**Tab F:** CUT&RUN antibodies (+ sequence stats)

*CUT&RUN sequence data [ncbi.nlm.nih.gov/geo/query/acc.cgi?acc=GSE249239]*

**Suppl. Table 2:** Compiled Captify data [Related **Figs** noted]

**Tab A: Fig. 1A -** Raw Data GST-BRD4-BD1 (EpiTriton Peptide Discovery Screen: 2 μM; Captify Peptide Discovery Screen: 1 nM & 30 nM)

**Tab B: Fig. 1B -** Raw Data GST-BRD4-BD1 (Signal-to-Background Comparison)

**Tab C: Fig. 1C -** Raw Data GST-BRD4-BD1 (Nuc Query and Discovery Screen)

**Tab D: Fig. 1D -** Raw Data GST-BRM-BD (Nuc Query and Discovery Screen)

**Tab E: Fig. 2A -** Raw Data GST-L3MBTL1-MBT (Peptide Query)

**Tab F: Fig. 2B -** Raw Data GST-L3MBTL1-MBT (Peptide Discovery Screen)

**Tab G: Fig. 2C -** Raw Data GST-L3MBTL1-MBT (Nuc Query and Discovery Screen)

**Tab H: Fig. 2D -** Raw Data GST-RAG2-PHD (Peptide Query and Discovery Screen)

**Tab I: Fig. 2E -** Raw Data GST-RAG2-PHD (Nuc Query and Discovery Screen)

**Tab J: Fig. 3A -** Raw Data GST-HP1β-CD (Nuc Query)

**Tab K: Fig. 3B -** Raw Data GST-ATRX-ADD (Nuc Query)

**Tab L: Fig. 3C -** Raw Data 6His-GLYR1-PWWP (salDNA Titration)

**Tab M: Fig. 3D -** Raw Data 6His-GLYR1-PWWP (Salt Optimization)

**Tab N: Fig. 3E -** Raw Data 6His-GLYR1-PWWP (Nuc Discovery Screen)

**Tab O: Fig. 3F -** Raw Data GST-CBX7-CD+ATL (Peptide Query)

**Tab P: Fig. 3G -** Raw Data GST-CBX7-CD+ATL (Nuc Query [-salDNA])

**Tab Q: Fig. 3H -** Raw Data GST-CBX7-CD+ATL (Nuc Query [+salDNA])

**Tab R: Fig. 3I -** Raw Data GST-CBX7-CD+ATL (Nuc Discovery Screen)

**Tab S: Fig. 4B -** Raw Data GST-HP1β-CD, GST-TAF3-PHD, GST-CBX7-CD+ATL (Luminex K-MetStat Nuc. Panel)

**Tab T: Fig. 5A -** Raw Data CUT&RUN Spike-In Nuc Barcode Results

**Tab U: Suppl. Fig. 1B -** BRD4-BD1 Comparison (TR-FRET and Captify-Alpha Peptide Query)

**Tab V: Suppl. Fig. 2A-C -** Raw Data (Peptide and Nuc Discovery Screens; Acyl Readers)

**Tab W: Suppl. Fig. 3A-H -** Raw Data (Peptide 2D Salt vs. Query Titrations; Acyl Readers)

**Tab X: Suppl. Fig. 4A-H -** Raw Data (Nuc 2D Salt vs. Query Titrations; Acyl Readers)

**Tab Y: Suppl. Fig. 7A-C -** Raw Data (Peptide and Nuc Discovery Screens; Methyl Readers)

**Tab Z: Suppl. Fig. 8A-G -** Raw Data (Peptide 2D Salt vs. Query Titrations; Methyl Readers)

**Tab AA: Suppl. Fig. 9A-G -** Raw Data (Nucl. 2D Salt vs. Query Titrations; Methyl Readers)

**Tab AB: Suppl. Fig. 10C-F -** Raw Data (Antibody and Reader MLA Binding)

**Tab AC: Suppl. Fig. 11A-F -** Raw Data GST-CBX7 CD & GST-CBX7 CD(F11A)-ATL (salDNA Titrations)

**Tab AD: Suppl. Fig. 12A-F** - Raw Data GST-CBX7 CD & GST-CBX7 CD(F11A)-ATL (Peptide & Nuc Queries)

**Tab AE: Suppl. Fig. 13A-B -** Raw Data (Chromatibody Acidic Patch Interactions)

### SUPPLEMENTARY FIGURE LEGENDS

**Suppl. Fig. 1.** **(A)** The assay previously known as dCypher™ is now named Captify™, with no distinction in how the assay is performed or its capabilities. The Captify-Alpha platform is a proximity assay combining biotinylated PTM-defined histone peptides or nucleosomes (the Targets), epitope-tagged reader proteins (the Queries) and the chemiluminescent Alpha System (streptavidin Donor and anti-tag Acceptor beads). Donor and Acceptor beads are brought into proximity by [Target : Query] engagement. Laser excitation (680 nm) of the Donor releases singlet oxygen that causes emission (520–620 nm) in proximal (within 200 nm) Donor beads; this luminescent signal is directly proportional to the amount of [Donor-Acceptor] bridged by the [Target : Query] interaction (see **Methods**). Binding is quantified by plotting Alpha Counts (fluorescence) as a function of protein concentration and expressed as relative EC50 [30] ($EC_{50}^{rel}$: see **Suppl. Table 2** for all from this study). **(B)** Histone peptide binding data generated using 6His-BRD4 BD1 (50 nM) in TR-FRET (1) and GST-BRD4 BD1 (1 nM) in Captify-Alpha (this study). **(C)** TR-FRET and Captify-Alpha deliver an almost identical ranking of [histone peptide : BRD4-BD1] interactions (1-10; highest to lowest assay signal for each approach).

**Suppl. Fig. 2. Lysine acyl readers bind a restricted range of PTM targets on nucleosomes *vs*. peptides.** Compiled Captify-Alpha discovery screen data for bromodomain (BD: BRD4, BRD4, BRM, BRG1) and YEATS domain (ENL, AF9, Gas41, YEATS2) Query (columns) binding to acylated histone peptide **(A)** and nucleosome **(B)** Targets (rows show selected interactions from all data in **Suppl. Table 2** [287 peptides and 77 nucleosomes]). Optimal conditions for each Query (*i.e.*, yielded maximal signal-to-background (S/B) within range $EC_{20}^{rel}$ – $EC_{80}^{rel}$ : **Suppl. Figs 3** and **4**) were used for discovery screens. Signals are normalized to maximum assay signal for each Query. **(C)** Max(imum) and Min(imum) assay signals for each Query with histone peptides or nucleosomes. **Key:** ac, acetylation; bu, butyrylation; cr, crotonylation;

H3.1NΔ32, nucleosome with H3 tail delete (residues 1-32); H4NΔ15, nucleosome with H4 tail delete (residues 1-15).

**Suppl. Fig. 3. Optimization of Captify-Alpha binding conditions for potential lysine acyl readers to peptide Targets.** 2D cross-titrations were used to optimize assay conditions for each Query **(A-H)** as maximal S/B from negative (N) (unmodified for background levels) and positive (P) control Target peptides. Each Query (nM; Y-axis) was cross-titrated against salt (NaCl, mM; X-axis) while measuring binding to histone peptides (100 nM). Assay signal for each Query heatmap is normalized to percent maximum signal (value noted). Follow-up discovery screens (*e.g.*, **Suppl. Figs 2A** & **C**) for each Query were generally performed at maximal S/B within range $EC_{20}^{rel}$ – $EC_{80}^{rel}$.

**Suppl. Fig. 4. Optimization of Captify-Alpha binding conditions for potential lysine acyl readers to nucleosome Targets.** 2D cross-titrations were used to optimize assay conditions for each Query **(A-H)** as maximal S/B from a predicted Target and unmodified nucleosome (Positive and Negative (background) controls respectively). Each Query (nM; Y-axis) was cross-titrated against salt (NaCl, mM; X-axis) while measuring binding to nucleosomes (10 nM). Assay signal for each Query heatmap is normalized to percent maximum signal (value noted). Follow-up discovery screens (*e.g.*, **Suppl. Figs 2B** & **C**) for each Query were performed at maximal S/B within range $EC_{20}^{rel}$ – $EC_{80}^{rel}$.

**Suppl. Fig. 5. Epitope tagged reader protein (Query) QC.** Queries were recombinantly expressed, purified, and quality estimated after SDS-PAGE and Coomassie staining. Material was used to generate data in **Fig. 1** and **Suppl. Figs 2 - 4**.

**Suppl. Fig. 6.** Native Top-Down Mass Spectrometry (nTDMS) was used to examine a potential [GST-RAG2-PHD : H3K4me3 nucleosome] complex (**Fig. 2F-G**). **(A)** After mixing GST-RAG2 PHD (5 μM) and H3K4me3 nucleosome (1 μM), nTDMS detected average mass of a complex (*i.e.* $MS^1$) corresponding to dimerized reader and nucleosome. Charge states 34+ and 35+ were isolated and activated by collisions with nitrogen to eject histone monomers. **(B)** The resulting $MS^2$ spectrum confirmed each histone including the H3K4me3 proteoform (shown as critical for complex formation by Captify: **Fig. 2E**).

**Suppl. Fig. 7. Lysine methyl readers bind a restricted range of PTM targets on nucleosomes *vs*. peptides.** Compiled Captify-Alpha discovery screening data for a range of Queries (columns) binding to histone peptide **(A)** and nucleosome **(B)** Targets (rows show selected interactions from all data in **Suppl. Table 2** [287 peptides and 77 nucleosomes]). Optimal conditions for each Query (*i.e.*, yielded maximal S/B within range $EC_{20}^{rel}$ – $EC_{80}^{rel}$ : **Suppl. Figs 8 - 10**) were used for discovery screens. Signals for each Query are normalized to maximum assay signal. **(C)** The Max and Min assay signals for each Query with histone peptides or nucleosomes (-, Not tested). **Note:** Extended data in **Fig. 2** (L3MBTL1 MBT, RAG2 PHD) and **Fig. 3** (HP1β CD, ATRX ADD, GLYR1 PWWP, CBX7 CD, CBX7 CD-ATL).

**Suppl. Fig. 8. Optimization of Captify-Alpha binding conditions for potential lysine methyl readers to peptide Targets.** 2D cross-titrations were used to optimize assay conditions for each Query **(A-G)** as maximal S/B from negative (N) (unmodified for background levels) and positive (P) control Target peptides. Each Query (nM; Y-axis) was cross-titrated against salt (NaCl, mM; X-axis) while measuring binding to histone peptides (100 nM). Assay signal for each Query heatmap is normalized to percent maximum signal (value noted). Follow-up discovery screens (*e.g.*, **Suppl. Fig 7A** & **C**) for each Query were performed at maximal S/B within range $EC_{20}^{rel}$ – $EC_{80}^{rel}$.

**Suppl. Fig. 9. Optimization of Captify-Alpha binding conditions for potential lysine methyl readers to nucleosome Targets.** 2D cross-titrations were used to optimize assay conditions for each Query **(A-G)** as maximal S/B from a predicted Target and unmodified nucleosome (Positive and Negative (background) controls respectively). Each Query (nM; Y-axis) was cross-titrated against salt (NaCl, mM; X-axis) while measuring binding to nucleosomes (10 nM). Assay signal for each Query heatmap is normalized to percent maximum signal (value as noted). Screening buffers for 6His-GLYR1-PWWP **(G)** were supplemented with 120 ng/mL salDNA to mitigate non-specific DNA binding. Follow-up discovery screens (*e.g.*, **Suppl. Fig. 7B** & **C**) for each Query were performed at maximal S/B within range $EC_{20}^{rel}$ – $EC_{80}^{rel}$.

**Suppl. Fig. 10. Unpredictable impact of the methyl lysine analog (MLA) when comparing antibody and reader binding to ([Kme3]) and ([K$_C$me3]).** **(A-B)** Structural comparison of Kme3 and K$_C$me3 (MLA) residues. **(C)** anti-H3K4me3 (*RevMab* 31-1039-00) binds equivalently to ([K4me3]) and ([K4$_C$me3]) nucleosomes. **(D)** GST RAG2-PHD binds equivalently to ([K4me3]) and ([K4$_C$me3]) nucleosomes. **(E)** anti-H3K9me3 (*ThermoFisher* 701784) binds equivalently to ([K9me3]) and ([K9$_C$me3]) nucleosomes. **(F)** GST ATRX ADD shows a profound preference for ([K9me3]) over ([K9$_C$me3]) nucleosomes (2,3).

**Suppl. Fig. 11. Impact of competitor DNA on nucleosome binding by CBX7 CD-ATL.** **(A)** salDNA competitor has no impact on GST CBX7-CD selectivity for ([H3K27me3]) over unmodified nucleosomes. **(B-C)** Presence of ATL (GST-CBX7 CD-ATL) increases background binding to unmodified nucleosomes but selectivity for ([H3K27me3]) is revealed by salDNA competitor (maximal S/B as noted) and largely lost by mutation of an aromatic cage residue within the chromodomain (GST CBX7 CD(F11A)-ATL : see also **Suppl. Fig. 12**).

**Suppl. Fig. 12. Nucleosome context is required for accurate determination of multivalent reader engagement by CBX7 CD-ATL. (A)** Titration of GST-CBX7 CD to histone peptides ($H3_{[1-20]}$ or $H3_{[15-34]}$) identifies equivalent binding to [H3K9me3 and H3K27me3] > [H3K9me2 and H3K27me2] (for impact of the ATL in this context compare to **Fig. 3F**). **(B)** Titration of GST-CBX7 CD(F11A)-ATL to histone peptides shows the chromodomain to mediate lysine-methyl binding (abolished by aromatic cage mutation F11A). **(C,D)** GST-CBX7 CD specifically binds ([H3K27me3]) in the nucleosome context and is unimpacted by salDNA competitor (up to 10 µg/mL) (compare to panel **A**, **Fig. 3G-H** and **Suppl. Fig. 11**). **(E,F)** Titration of GST-CBX7 CD(F11A)-ATL shows the ATL mediates non-specific nucleosome binding (compare to **Fig 3G-H** and **Suppl. Fig. 11**).

**Suppl. Fig. 13**. **Increased assay stringency reveals loss-of-function interactions with the nucleosome acid-patch.** H2BK120ub interferes with the binding of multiple proteins to the nucleosome acidic patch (4-6). **(A-B)** The anti-acid-patch chromatibody VHH (7) is unimpacted by H2BK120ub (binding relative to unmodified nucleosome) in no-wash Captify-Alpha but shows ~10-fold reduced binding in the increased stringency washable format Captify -Luminex. Controls are H2AK119ub (distal to the acidic patch (8)) and acid-patch mutant H2AE92K (9).

**Suppl. Fig. 14. Epitope tagged reader protein (Query) QC.** Queries were recombinantly expressed, purified, and quality estimated after SDS-PAGE and Coomassie staining. Material was used to generate data in **Figs 2 - 5** and **Suppl. Figs 7 - 13**.

**Suppl. Fig. 15. Readers can deliver antibody-like PTM profiling in CUT&RUN genomic mapping. (A)** Heat map of GST-TAF3 PHD and H3K4me3 CUT&RUN reads aligned to the transcription start site (TSS ±2kb) of 19,292 genes in K562 cells. Rows were k-means clustered using the ChAsE chromatin analysis tool (10). IgG and $\alpha$GST inform on assay background. **(B)**

H3K4me3 (and GST TAF3-PHD) are most associated with gene promoters for the most highly abundant mRNAs. CUT&RUN reads were aligned to protein coding genes (normalized for length), spanning TSS to transcription termination site (TSS-TTS; ±2kb), and binned by decile (color code in legend) according to K562 RNA-seq expression data (as in **Methods**).

### SUPPLEMENTAL REFERENCES


1. Jung, M., Philpott, M., Muller, S., Schulze, J., Badock, V., Eberspacher, U., Moosmayer, D., Bader, B., Schmees, N., Fernandez-Montalvan, A. *et al.* (2014) Affinity map of bromodomain protein 4 (BRD4) interactions with the histone H4 tail and the small molecule inhibitor JQ1. *J Biol Chem*, **289**, 9304-9319.

2. Weinzapfel, E.N., Fedder-Semmes, K.N., Sun, Z.W. and Keogh, M.C. (2024) Beyond the tail: the consequence of context in histone post-translational modification and chromatin research. *Biochem J*, **481**, 219-244.

3. Seeliger, D., Soeroes, S., Klingberg, R., Schwarzer, D., Grubmuller, H. and Fischle, W. (2012) Quantitative assessment of protein interaction with methyl-lysine analogues by hybrid computational and experimental approaches. *ACS Chem Biol*, **7**, 150-154.

4. Barbera, A.J., Chodaparambil, J.V., Kelley-Clarke, B., Joukov, V., Walter, J.C., Luger, K. and Kaye, K.M. (2006) The nucleosomal surface as a docking station for Kaposi's sarcoma herpesvirus LANA. *Science*, **311**, 856-861.

5. Makde, R.D., England, J.R., Yennawar, H.P. and Tan, S. (2010) Structure of RCC1 chromatin factor bound to the nucleosome core particle. *Nature*, **467**, 562-566.

6. Hicks, C.W., Rahman, S., Gloor, S.L., Fields, J.K., Husby, N.L., Vaidya, A., Maier, K.E., Morgan, M., Keogh, M.C. and Wolberger, C. (2024) Ubiquitinated histone H2B as gatekeeper of the nucleosome acidic patch. *Nucleic Acids Res*, **52**, 9978-9995.

7. Jullien, D., Vignard, J., Fedor, Y., Bery, N., Olichon, A., Crozatier, M., Erard, M., Cassard, H., Ducommun, B., Salles, B. *et al.* (2016) Chromatibody, a novel non-invasive molecular tool to explore and manipulate chromatin in living cells. *J Cell Sci*, **129**, 2673-2683.

8. Thomas, J.F., Valencia-Sanchez, M.I., Tamburri, S., Gloor, S.L., Rustichelli, S., Godinez-Lopez, V., De Ioannes, P., Lee, R., Abini-Agbomson, S., Gretarsson, K. *et al.* (2023) Structural basis of histone H2A lysine 119 deubiquitination by Polycomb Repressive Deubiquitinase BAP1/ASXL1. *Sci Adv.*, **9(32):eadg9832**.

9. Kalashnikova, A.A., Porter-Goff, M.E., Muthurajan, U.M., Luger, K. and Hansen, J.C. (2013) The role of the nucleosome acidic patch in modulating higher order chromatin structure. *J R Soc Interface*, **10**, 20121022.

10. Younesy, H., Nielsen, C.B., Lorincz, M.C., Jones, S.J., Karimi, M.M. and Moller, T. (2016) ChAsE: chromatin analysis and exploration tool. *Bioinformatics*, **32**, 3324-3326.
