## Supplemental Figures (1-15) for "Nucleosome context regulates chromatin reader preference"

**Suppl. Fig 1_Marunde et al**

A

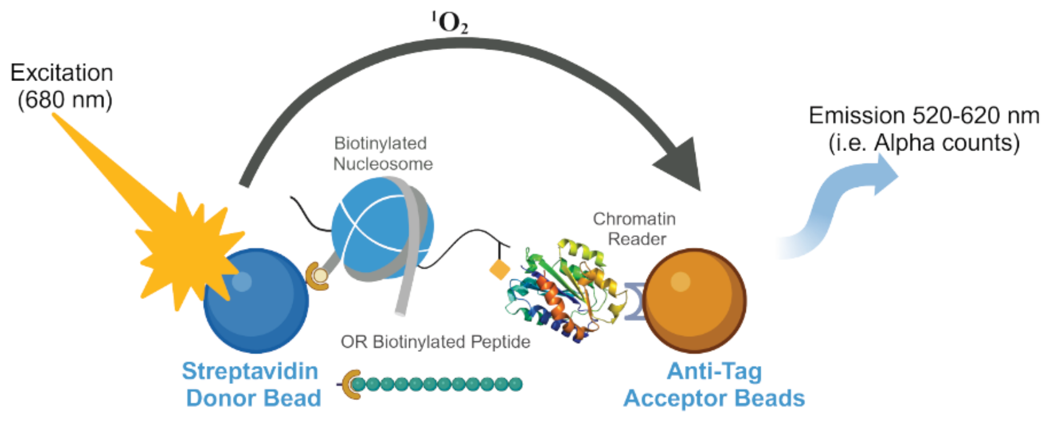


B

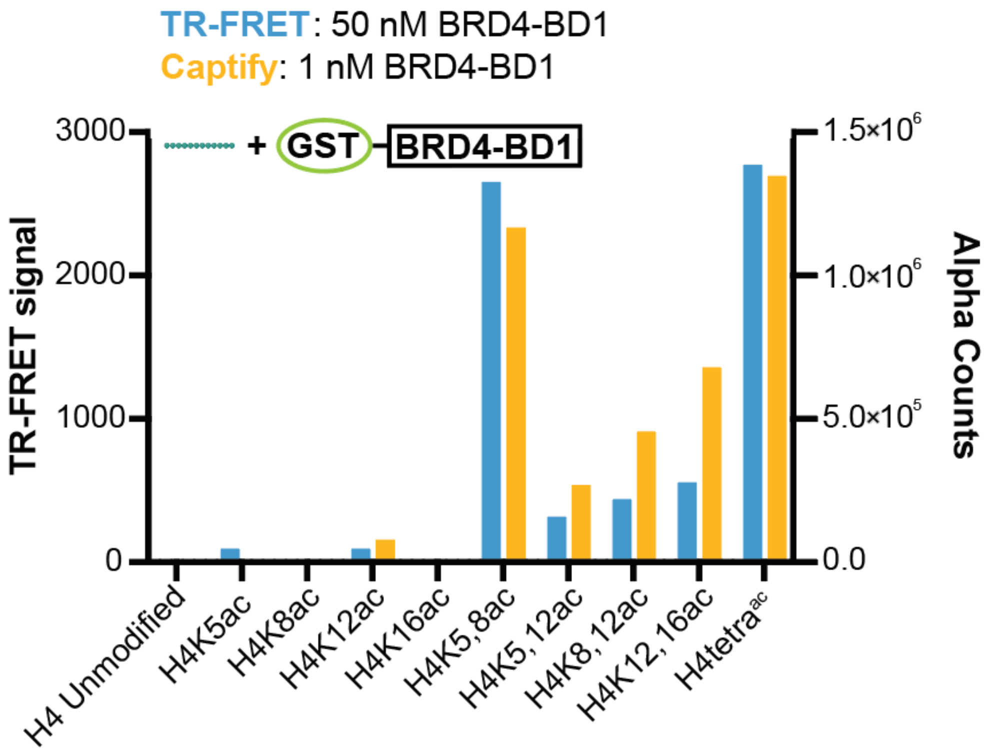


C

| Peptide TARGET | TR-FRET Rank | Captify Rank |
|---|---|---|
| H4tetra$^{ac}$ | 1 | 1 |
| H4K5,8ac | 2 | 2 |
| H4K12,16ac | 3 | 3 |
| H4K8,12ac | 4 | 4 |
| H4K5,12ac | 5 | 5 |
| H4K5ac | 6 | 7 |
| H4K12ac | 7 | 6 |
| H4K8ac | 8 | 8 |
| H4K16ac | 9 | 9 |
| H4 Unmodified | 10 | 10 |

Suppl. Fig 2_Marunde et al

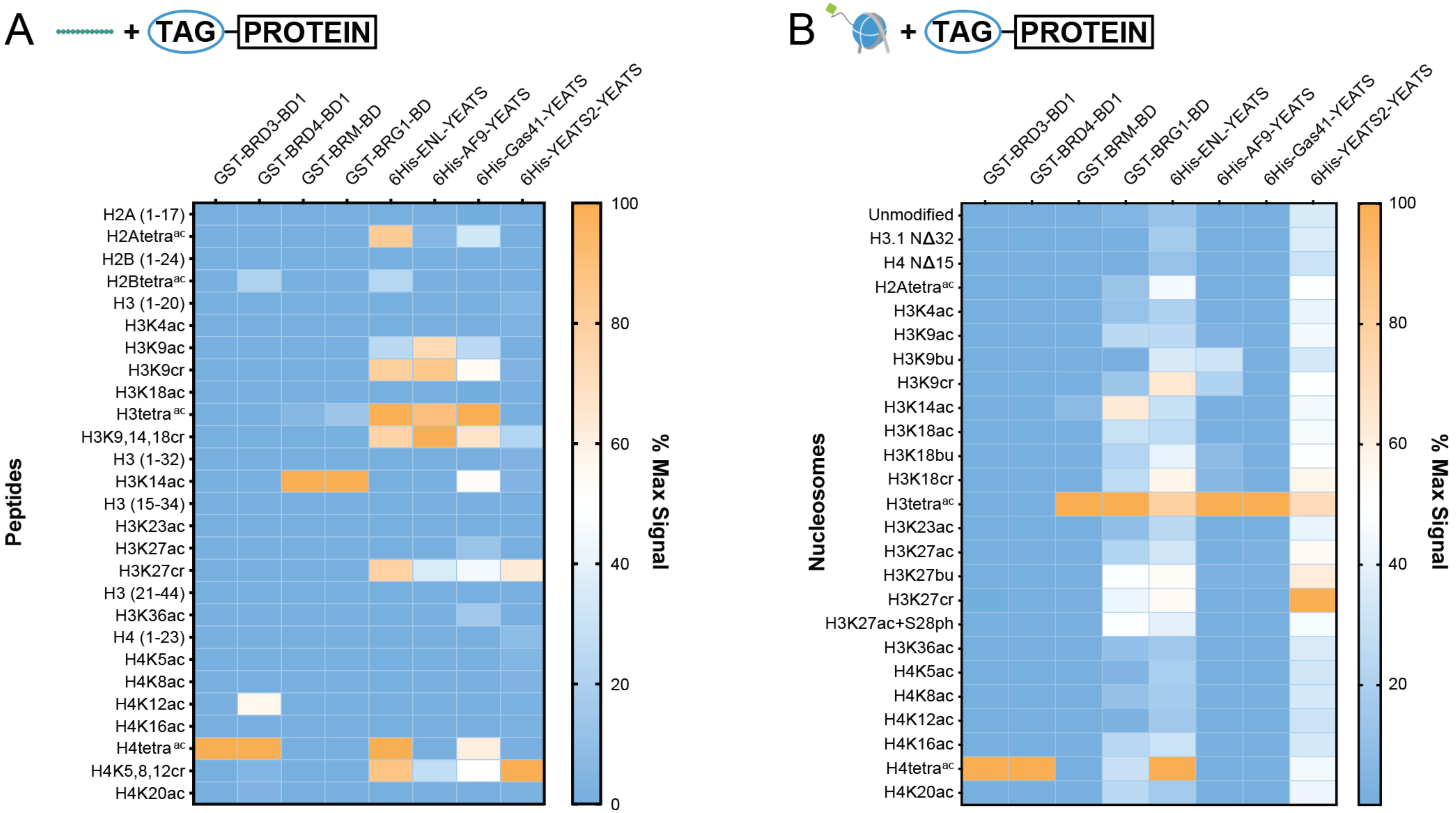


C

| TARGET (Signal) | BRD3 | BRD4 | BRM | BRG1 | ENL | AF9 | Gas41 | YEATS2 |
|---|---|---|---|---|---|---|---|---|
| Peptide (Max) | 107,992 | 557,686 | 189,845 | 361,785 | 355,312 | 213,795 | 316,755 | 82,558 |
| Peptide (Min) | 356 | 133 | 257 | 112 | 163 | 10 | 0 | 264 |
| Nuc (Max) | 316,640 | 134,981 | 107,109 | 494,504 | 195,874 | 75,244 | 30,298 | 4,796 |
| Nuc (Min) | 307 | 685 | 688 | 4773 | 23,548 | 96 | 139 | 1,451 |

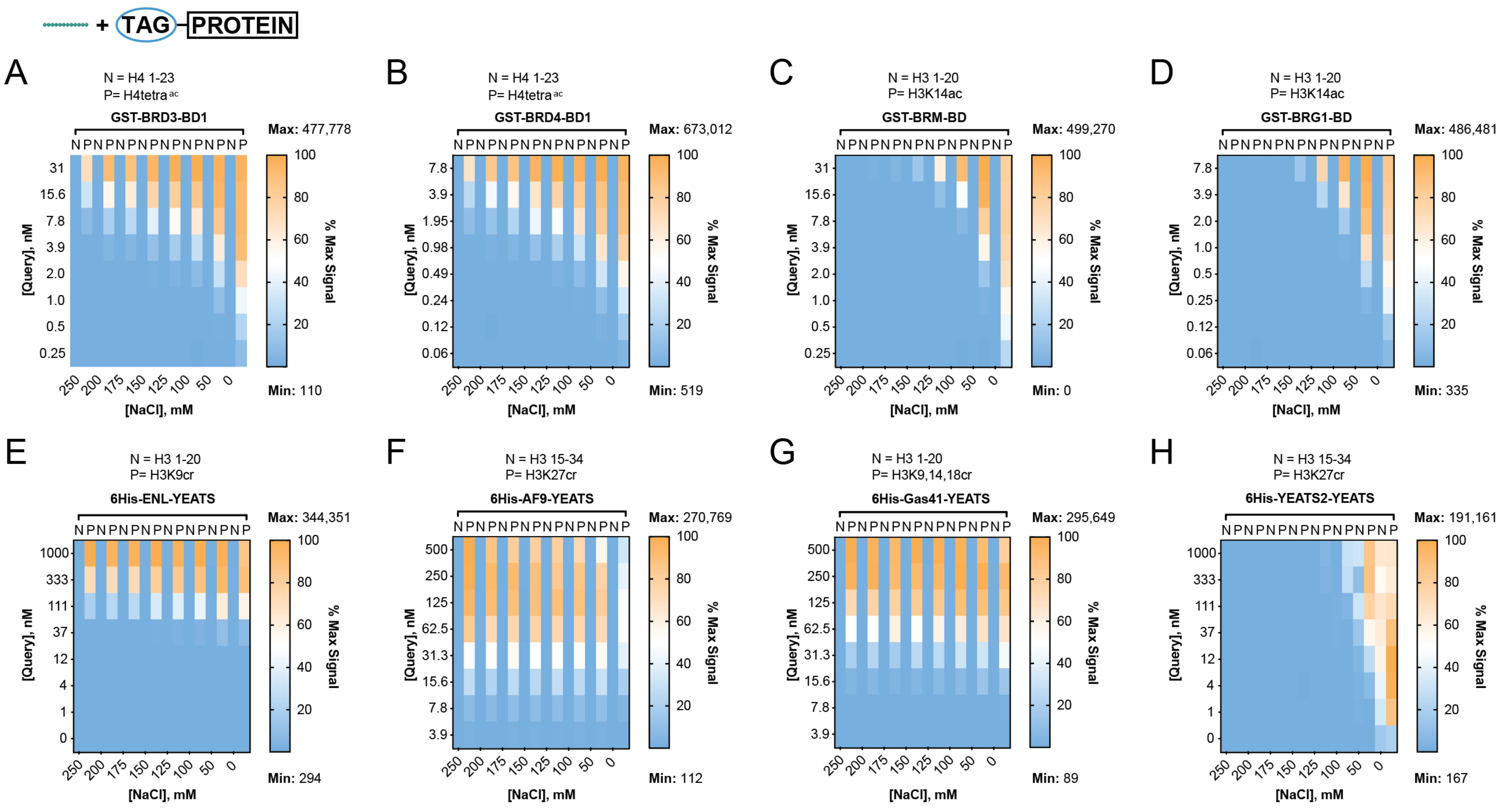
TAG
PROTEIN
A
N = H4 1-23
P= H4tetra ac
GST-BRD3-BD1
Max: 477,778
Min: 110
B
N = H4 1-23
P= H4tetra ac
GST-BRD4-BD1
Max: 673,012
Min: 519
C
N = H3 1-20
P= H3K14ac
GST-BRM-BD
Max: 499,270
Min: 0
D
N = H3 1-20
P= H3K14ac
GST-BRG1-BD
Max: 486,481
Min: 335
E
N = H3 1-20
P= H3K9cr
6His-ENL-YEATS
Max: 344,351
Min: 294
F
N = H3 15-34
P= H3K27cr
6His-AF9-YEATS
Max: 270,769
Min: 112
G
N = H3 1-20
P= H3K9,14,18cr
6His-Gas41-YEATS
Max: 295,649
Min: 89
H
N = H3 15-34
P= H3K27cr
6His-YEATS2-YEATS
Max: 191,161
Min: 167
[Query], nM
[NaCl], mM
% Max Signal

Suppl. Fig 4_Marunde et al

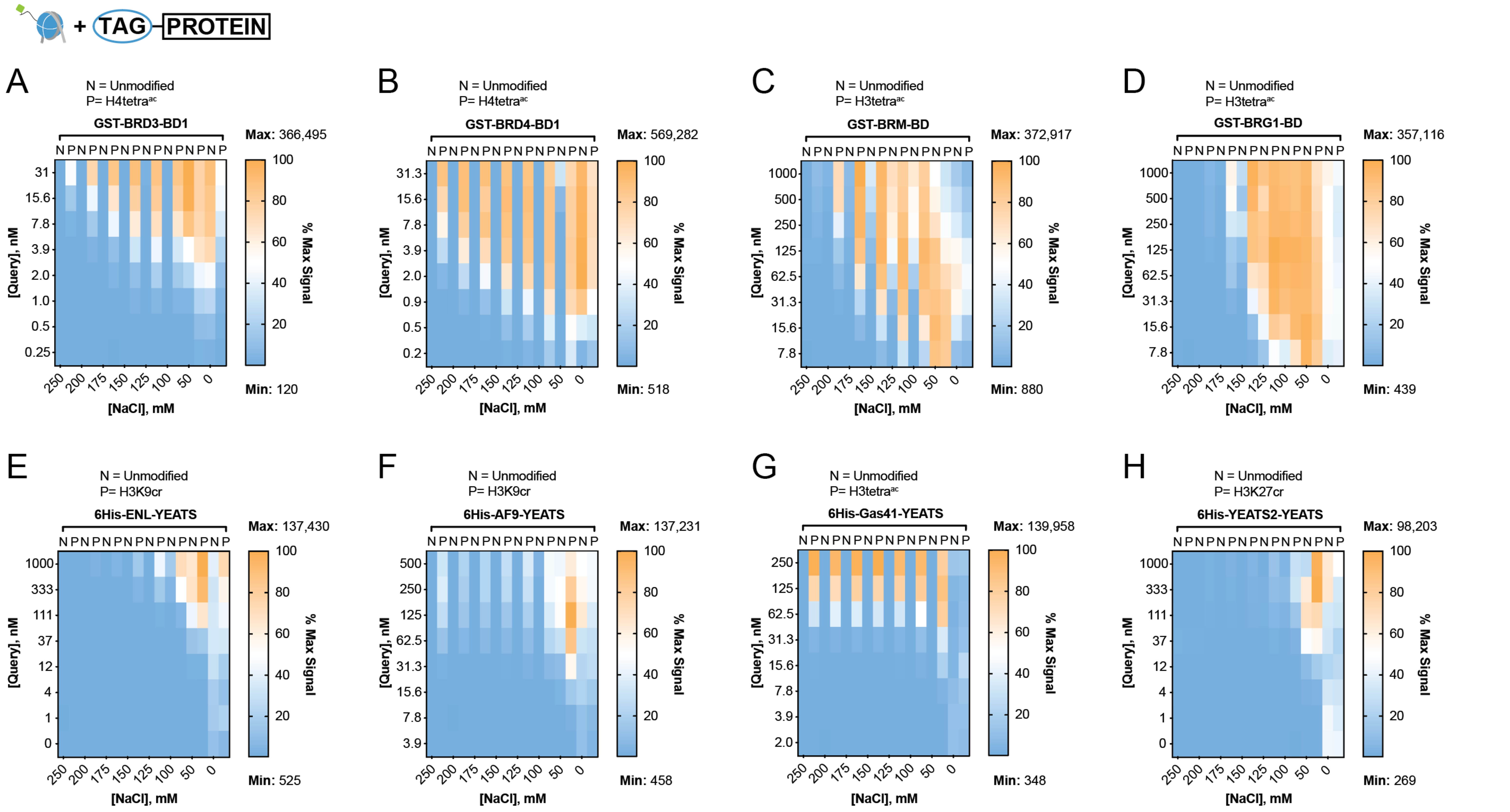

Suppl. Fig 5_Marunde et al

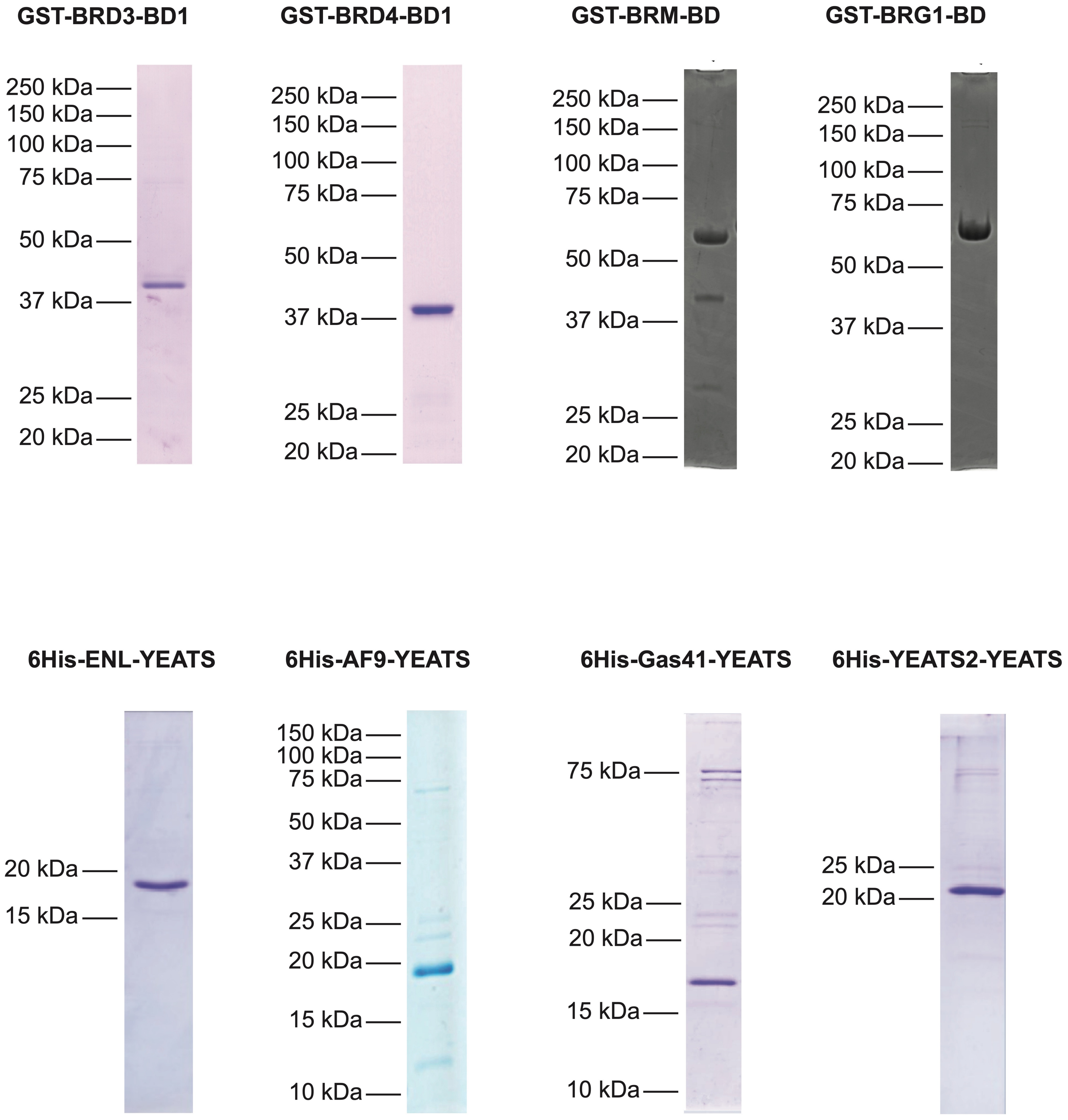

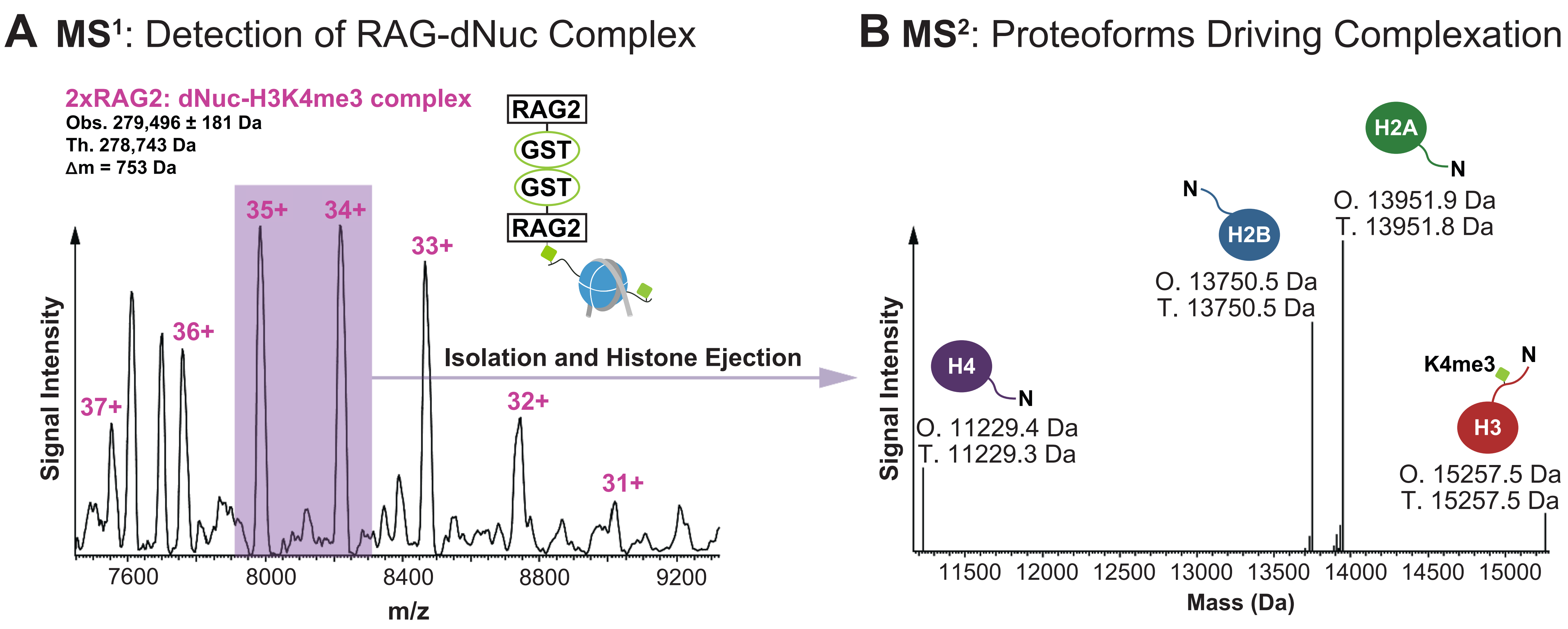
A MS¹: Detection of RAG-dNuc Complex
2xRAG2: dNuc-H3K4me3 complex
Obs. 279,496 ± 181 Da
Th. 278,743 Da
Δm = 753 Da
RAG2
GST
GST
RAG2
35+
34+
33+
36+
37+
32+
31+
Isolation and Histone Ejection
Signal Intensity
7600
8000
8400
8800
9200
m/z
B MS²: Proteoforms Driving Complexation
H2A
N
O. 13951.9 Da
T. 13951.8 Da
H2B
O. 13750.5 Da
T. 13750.5 Da
H4
O. 11229.4 Da
T. 11229.3 Da
K4me3
H3
O. 15257.5 Da
T. 15257.5 Da
Signal Intensity
11500
12000
12500
13000
13500
14000
14500
15000
Mass (Da)

**Suppl. Fig 7_Marunde et al**

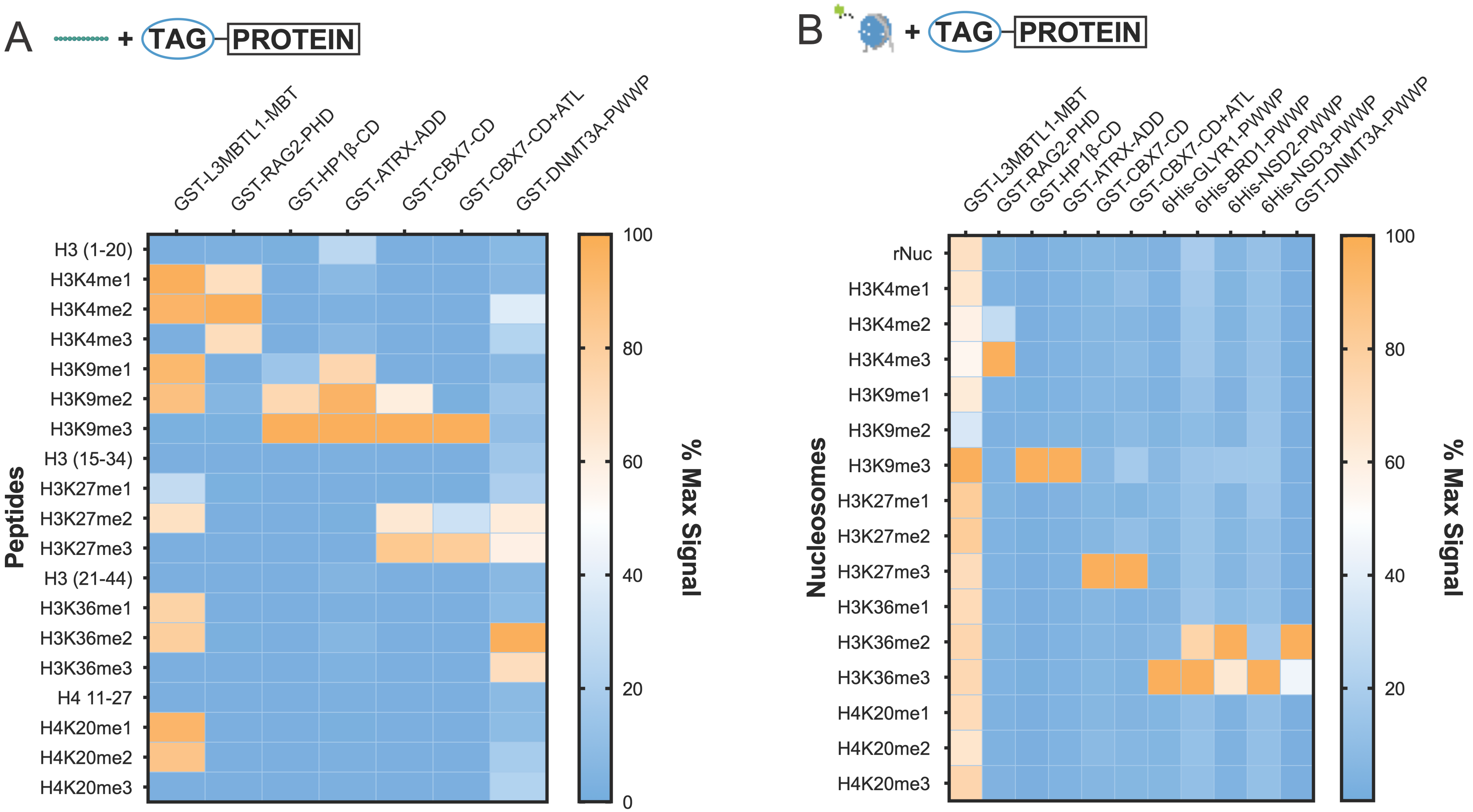


C

| TARGET (Signal) | L3MBTL1 | RAG2 | HP1β | ATRX | CBX7-CD | CBX7-CD+ATL | GLYR1 | BRD1 | NSD2 | NSD3 | DNMT3A |
|---|---|---|---|---|---|---|---|---|---|---|---|
| Peptide (Max) | 281,106 | 286,519 | 265,629 | 234,173 | 545,984 | 485,902 | - | - | - | - | 259,532 |
| Peptide (Min) | 33 | 436 | 343 | 621 | 0 | 0 | - | - | - | - | 13,965 |
| Nuc (Max) | 2,193 | 43,978 | 263,967 | 34,763 | 17,403 | 69,204 | 42,306 | 40,539 | 95,705 | 5,581 | 242,275 |
| Nuc (Min) | 774 | 621 | 547 | 684 | 622 | 1,674 | 542 | 3,066 | 1,591 | 504 | 186 |

Suppl. Fig 8_Marunde et al

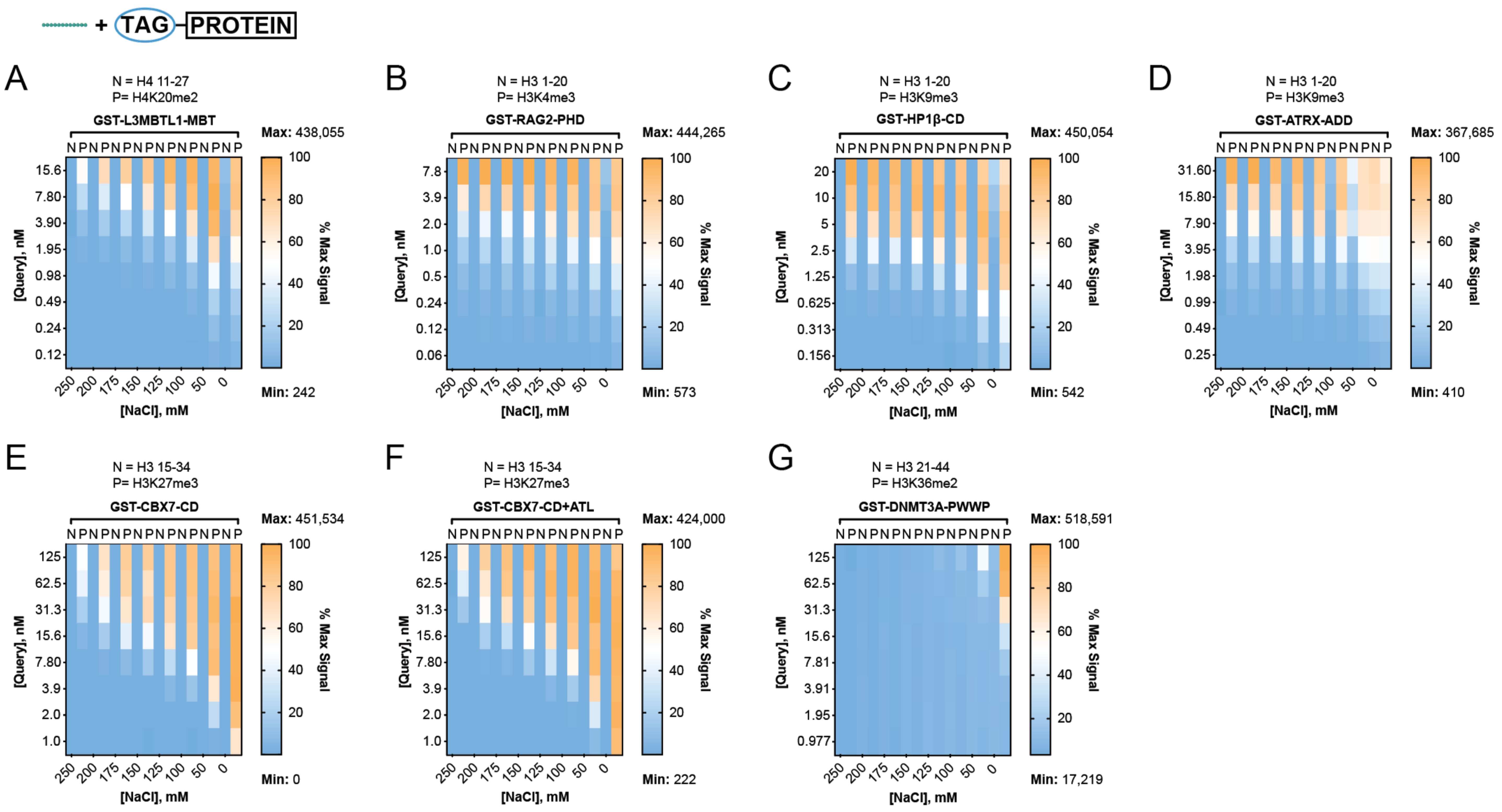

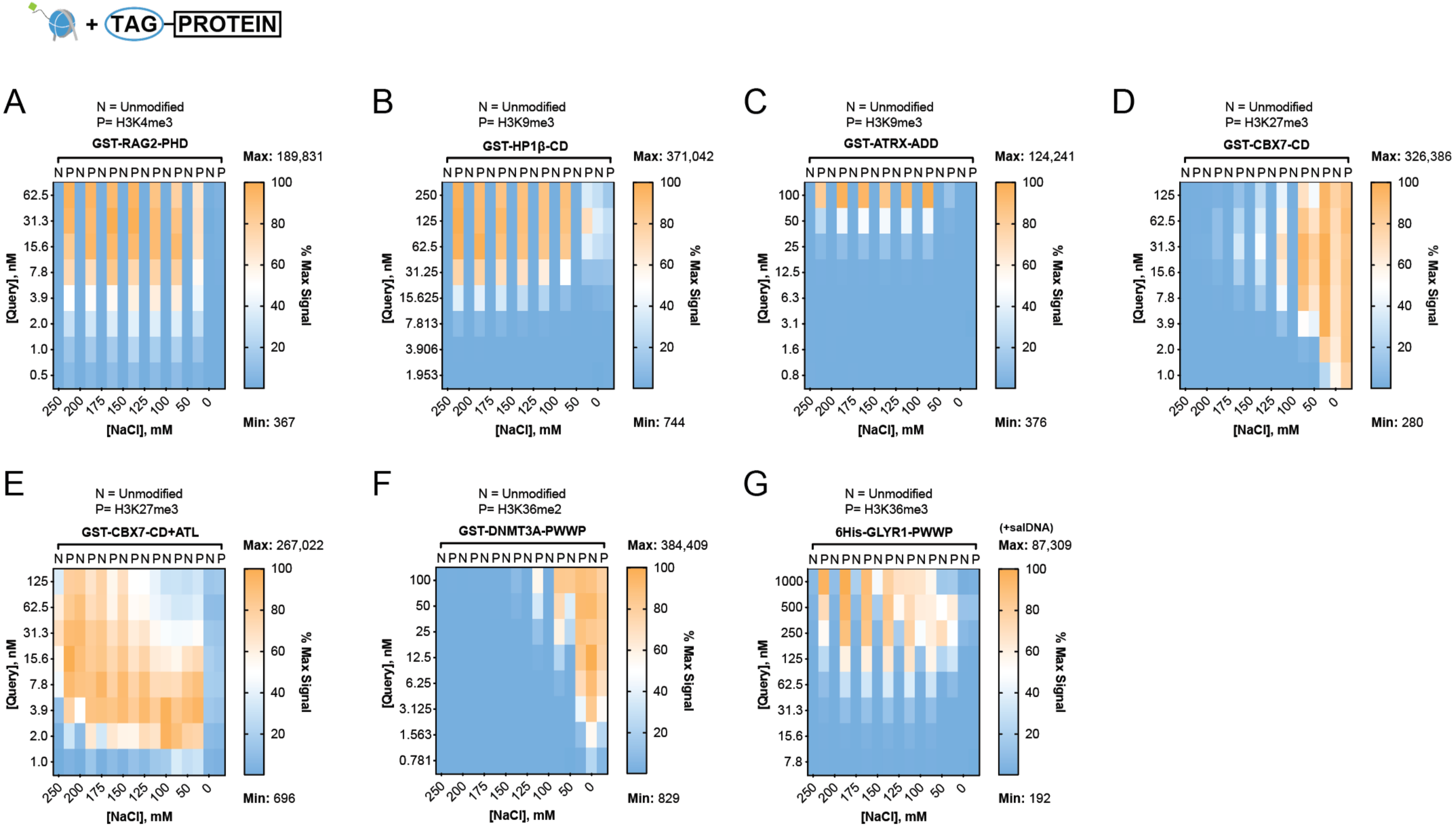

+ TAG PROTEIN
A
N = Unmodified
P= H3K4me3
GST-RAG2-PHD
Max: 189,831
Min: 367
B
N = Unmodified
P= H3K9me3
GST-HP1β-CD
Max: 371,042
Min: 744
C
N = Unmodified
P= H3K9me3
GST-ATRX-ADD
Max: 124,241
Min: 376
D
N = Unmodified
P= H3K27me3
GST-CBX7-CD
Max: 326,386
Min: 280
E
N = Unmodified
P= H3K27me3
GST-CBX7-CD+ATL
Max: 267,022
Min: 696
F
N = Unmodified
P= H3K36me2
GST-DNMT3A-PWWP
Max: 384,409
Min: 829
G
N = Unmodified
P= H3K36me3
6His-GLYR1-PWWP
(+salDNA)
Max: 87,309
Min: 192
[Query], nM
[NaCl], mM
% Max Signal

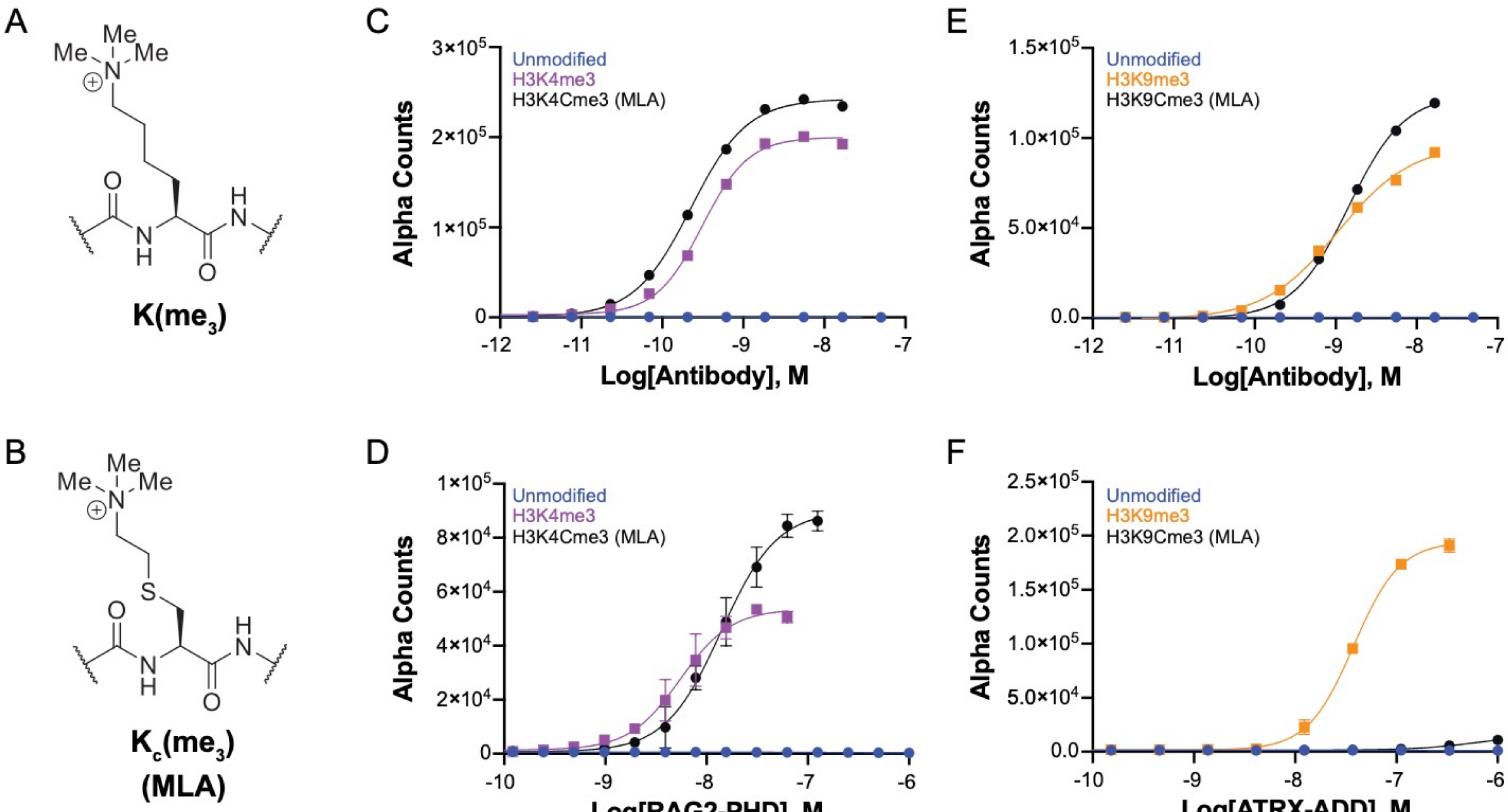
A
Me
Me
Me
N
O
H
N
N
H
O
K(me3)
B
Me
Me
Me
N
S
O
H
N
N
H
O
Kc(me3)
(MLA)
C
Unmodified
H3K4me3
H3K4Cme3 (MLA)
Alpha Counts
3×10^5
2×10^5
1×10^5
0
-12
-11
-10
-9
-8
-7
Log[Antibody], M
D
Unmodified
H3K4me3
H3K4Cme3 (MLA)
Alpha Counts
1×10^5
8×10^4
6×10^4
4×10^4
2×10^4
0
-10
-9
-8
-7
-6
Log[RAG2-PHD], M
E
Unmodified
H3K9me3
H3K9Cme3 (MLA)
Alpha Counts
1.5×10^5
1.0×10^5
5.0×10^4
0.0
-12
-11
-10
-9
-8
-7
Log[Antibody], M
F
Unmodified
H3K9me3
H3K9Cme3 (MLA)
Alpha Counts
2.5×10^5
2.0×10^5
1.5×10^5
1.0×10^5
5.0×10^4
0.0
-10
-9
-8
-7
-6
Log[ATRX-ADD], M

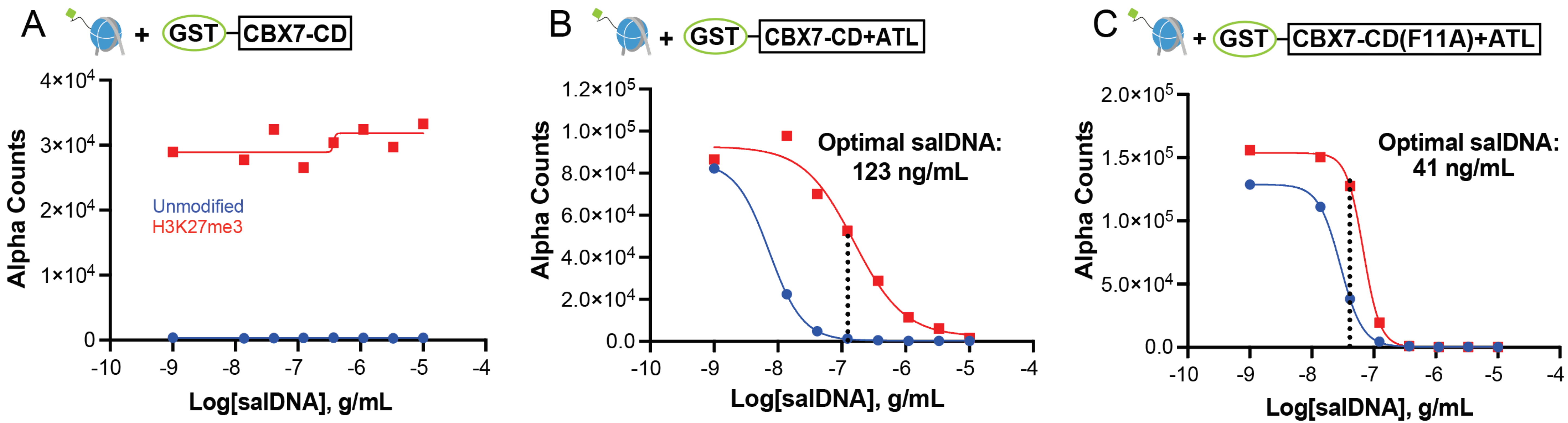
A
+ GST CBX7-CD
Alpha Counts
4×10^4
3×10^4
2×10^4
1×10^4
0
Unmodified
H3K27me3
-10 -9 -8 -7 -6 -5 -4
Log[salDNA], g/mL
B
+ GST CBX7-CD+ATL
Alpha Counts
1.2×10^5
1.0×10^5
8.0×10^4
6.0×10^4
4.0×10^4
2.0×10^4
0.0
Optimal salDNA:
123 ng/mL
-10 -9 -8 -7 -6 -5 -4
Log[salDNA], g/mL
C
+ GST CBX7-CD(F11A)+ATL
Alpha Counts
2.0×10^5
1.5×10^5
1.0×10^5
5.0×10^4
0.0
Optimal salDNA:
41 ng/mL
-10 -9 -8 -7 -6 -5 -4
Log[salDNA], g/mL

Suppl. Fig 12_Marunde et al

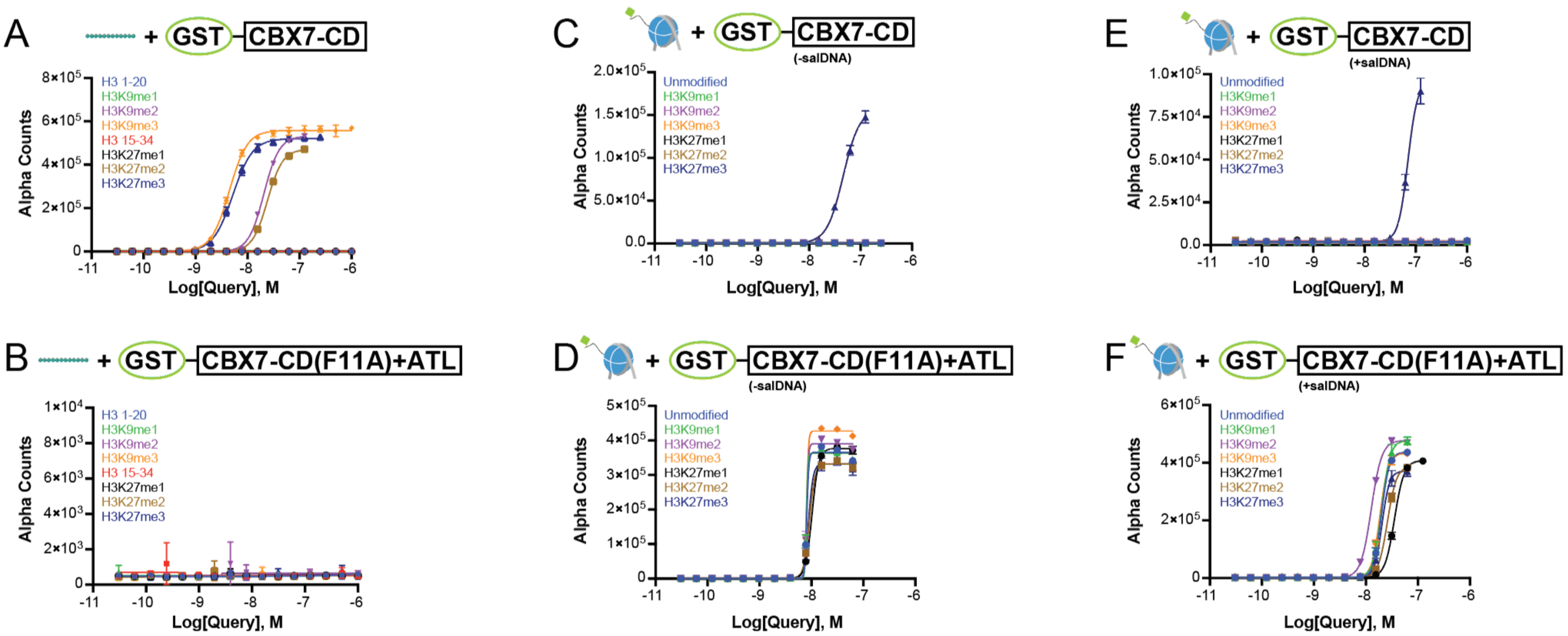

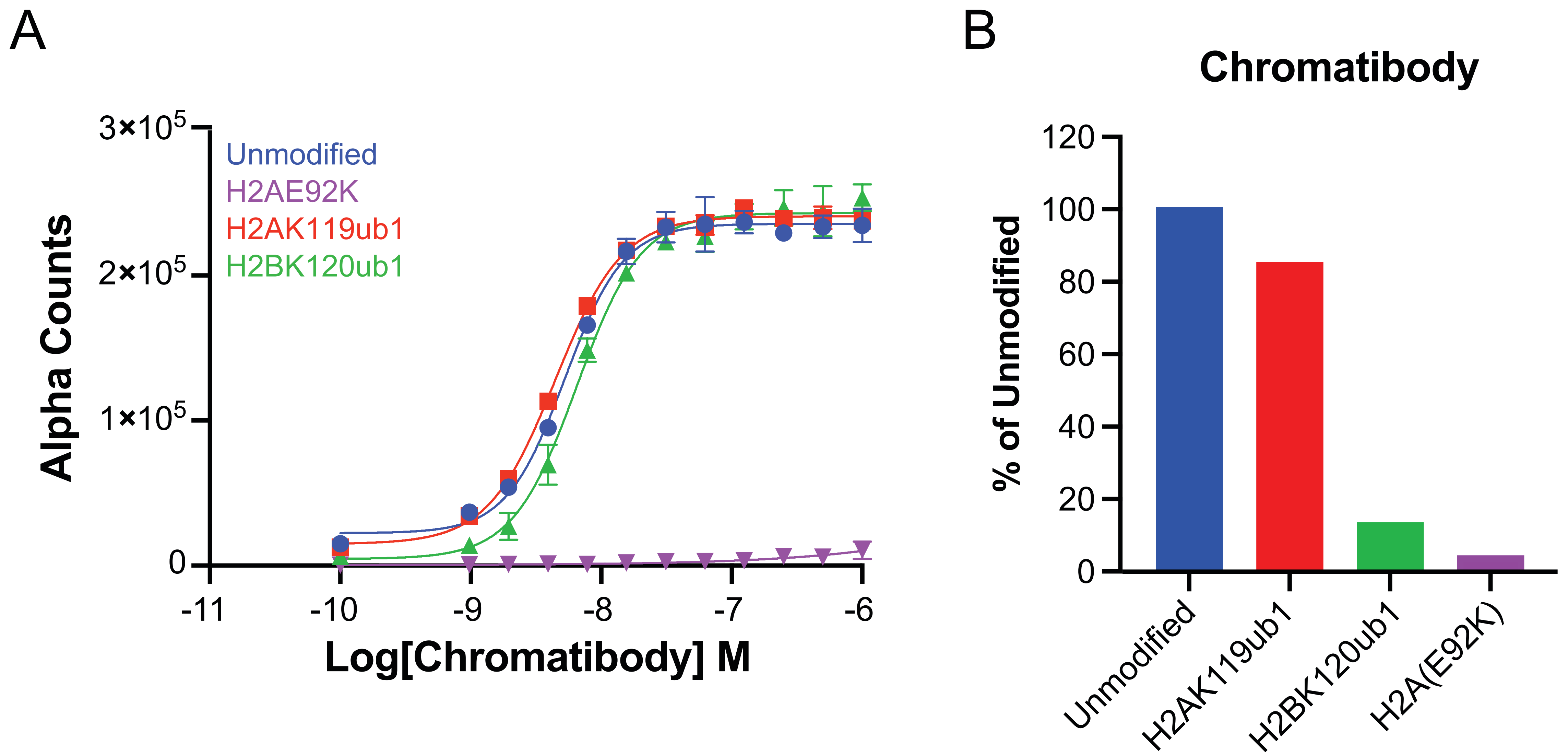
A
3×10^5
2×10^5
1×10^5
0
Unmodified
H2AE92K
H2AK119ub1
H2BK120ub1
Alpha Counts
-11
-10
-9
-8
-7
-6
Log[Chromatibody] M
B
Chromatibody
120
100
80
60
40
20
0
% of Unmodified
Unmodified
H2AK119ub1
H2BK120ub1
H2A(E92K)

Suppl. Fig 14_Marunde et al

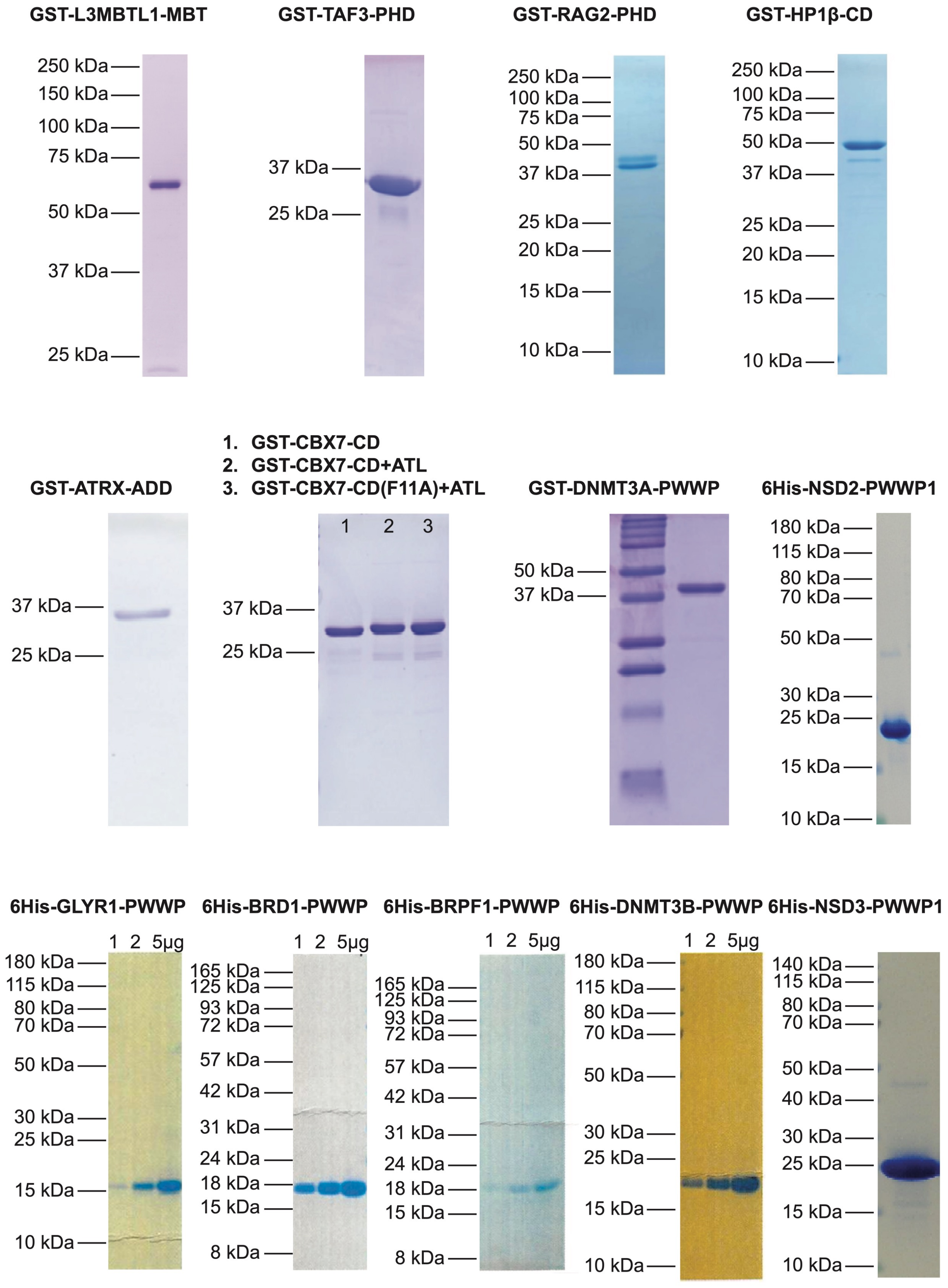

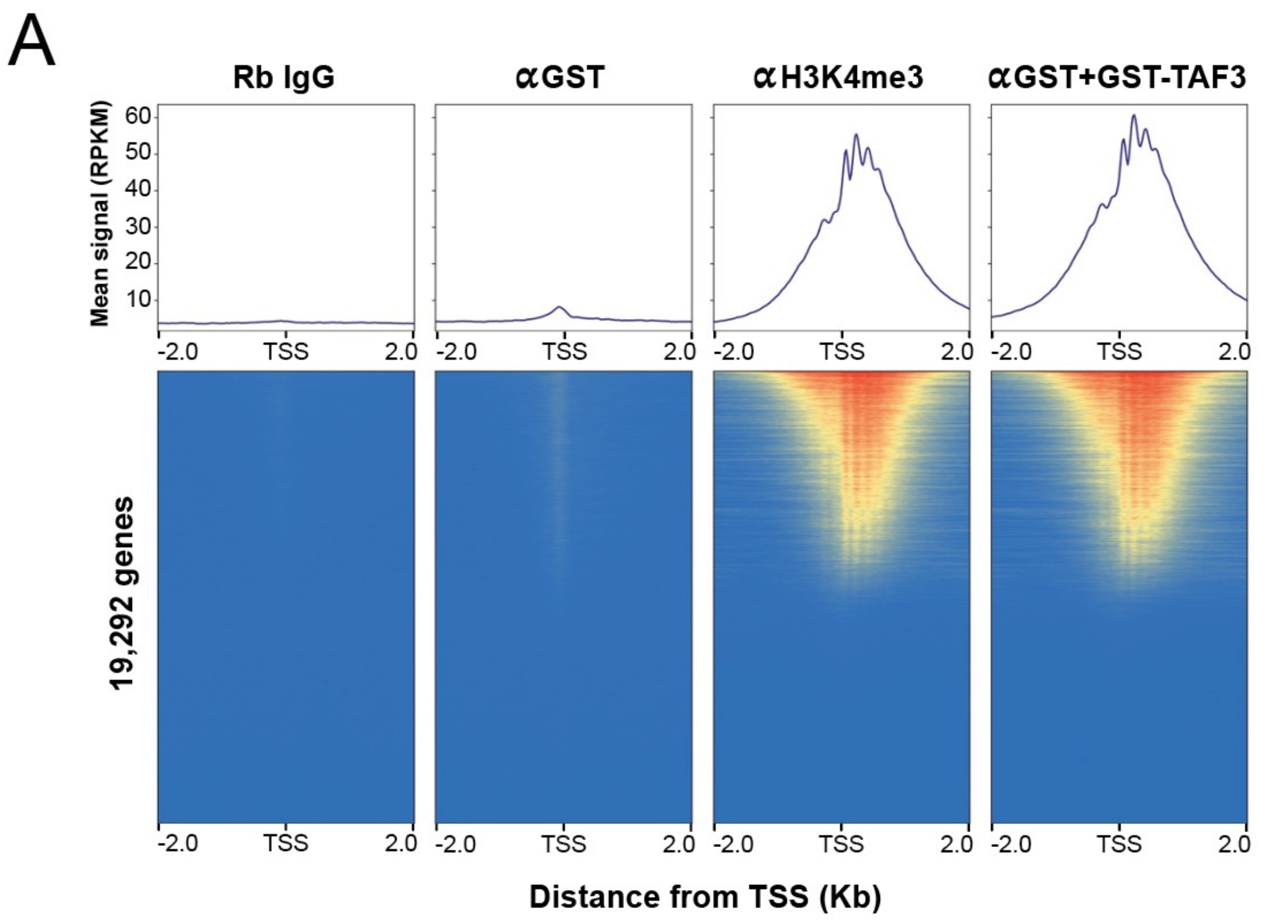
A
Rb IgG
αGST
αH3K4me3
αGST+GST-TAF3
Mean signal (RPKM)
60
50
40
30
20
10
-2.0
TSS
2.0
19,292 genes
Distance from TSS (Kb)

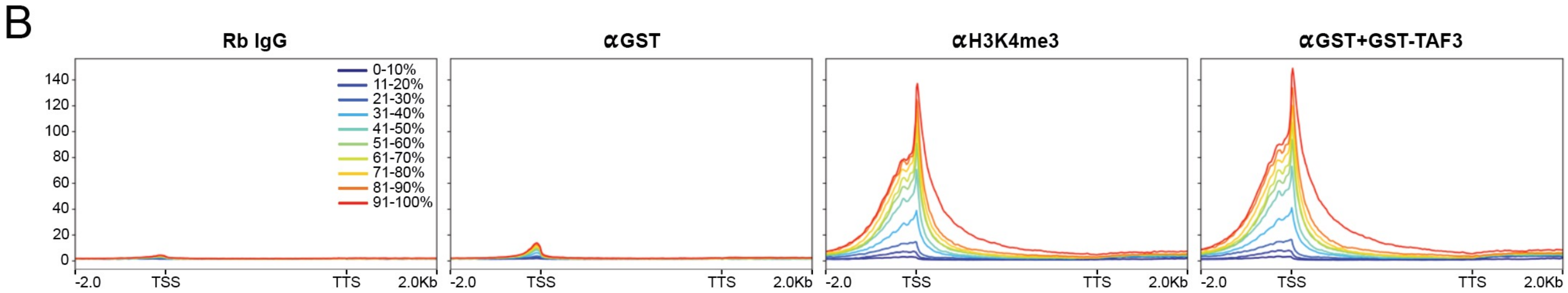
B
Rb IgG
αGST
αH3K4me3
αGST+GST-TAF3
0-10%
11-20%
21-30%
31-40%
41-50%
51-60%
61-70%
71-80%
81-90%
91-100%
140
120
100
80
60
40
20
0
-2.0
TSS
TTS
2.0Kb
